# Unicore Enables Scalable and Accurate Phylogenetic Reconstruction with Structural Core Genes

**DOI:** 10.1101/2024.12.22.629535

**Authors:** Dongwook Kim, Sukhwan Park, Martin Steinegger

## Abstract

The analysis of single-copy core genes, common to most members of a clade, is important for key tasks in biology including phylogenetic reconstruction and assessing genome quality. Core genes are traditionally identified by the analysis of amino acid similarities among proteomes, but can also be defined using structures, which bear potential in deep clades beyond the twilight zone of amino acids. Despite breakthroughs in accurate AI-driven protein structure prediction, obtaining full 3D structural models on a proteomic scale is still prohibitively slow. Here, we present Unicore, a novel method for identifying structural core genes at a scale suitable for downstream phylogenetic analysis. By applying the ProstT5 protein language model to the input proteomes to obtain their 3Di structural strings, Unicore saves over three orders of magnitude in runtime compared to a full 3D prediction. Using Foldseek clustering, Unicore identifies single-copy structures universally present in the species and aligns them using Foldmason. These structural core gene alignments are projected back to amino acid information for downstream phylogenetic analysis. We demonstrate that this approach defines core genes with linear run-time scaling over the number of species, up to 56 times faster than OrthoFinder, while reconstructing phylogenetic relationships congruent with conventional approaches. Unicore is universally applicable to any given set of taxa, even spanning superkingdoms and overcoming limitations of previous methods requiring orthologs of fixed taxonomic scope, and is available as a free and open source software at https://github.com/steineggerlab/unicore.

## Introduction

Understanding the evolutionary relationships among organisms through phylogenetic analysis is a fundamental goal in biology. Phylogenomics allows us to reconstruct the tree of life, shedding light on how different species are related and how they have diverged over time (1). A common phylogenomic approach leverages single-copy core genes—universally conserved across taxa—as reliable markers for reconstructing phylogenetic relationships (2).

State-of-the-art core gene databases, including OrthoDB (3), OMA (4), UBCG (5) or UFCG (6), offer lists of single-copy core genes for specific clades. These gene sets are detected from genomes through sequence-based orthology inference (7) or profile hidden Markov model (HMM)-based homology searches (8). However, HMM- or all-vs-all search-based (9) approaches are computationally intensive and less effective for newly discovered clades, particularly when they are in the ‘twilight zone’ from the known clades, where amino acid similarity is too low to detect homology (10, 11). Protein structure-based methods, like Fold-seek (12) using the 3Di alphabet, overcome these challenges by detecting homologs below the twilight zone through structural searches (11, 13, 14). Furthermore, Foldseek’s linear-scaling time clustering enables rapid proteome-wide comparisons (15, 16).

The advent of AlphaFold2 (17) has opened the door for much broader use of structural information in phylogenetic analysis by lifting the dependency on limitedly available experimentally determined structures (18, 19). Recent research shows that Foldseek’s 3Di alignments reveal deep evolutionary relationships and robust phylogenetic signals even for highly divergent sequences (20–22). However, applying AlphaFold2 or similar methods to even dozens of full proteomes for core gene detection would require months, limiting its use. Alternatively, protein language models (pLMs; (23–25)) can be used to extract structural features directly from primary amino acid sequences, eliminating the need for computationally intensive structure prediction methods. Among them, ProstT5 (26) stands out, which is a bilingual pLM trained on millions of sequence-structure pairs from the AFDB (27). ProstT5 rapidly converts amino acid sequences into 3Di strings 3,600-fold faster than AlphaFold2-ColabFold (28), thereby making structure-informed large-scale phylogenetic analysis feasible.

Here we present Unicore, a scalable and accurate method for structure-based core gene definition and phylogenetic inference. Unicore leverages predicted 3Di sequences from the ProstT5 model and linear-time comparison methods (16) to accelerate large-scale proteome analysis. It identifies single-copy ‘structural core genes’ on the fly without relying on pre-computed gene sets (Fig. 1). By integrating Foldseek’s rapid structural comparisons, Unicore identifies conserved structural genes across proteomes, generates structural alignments from 3Di strings using FoldMason (29), and infers phylogenetic trees with maximum likelihood methods, such as IQ-TREE (30), FastTree (31), and RAxML (32).

**Fig. 1.**
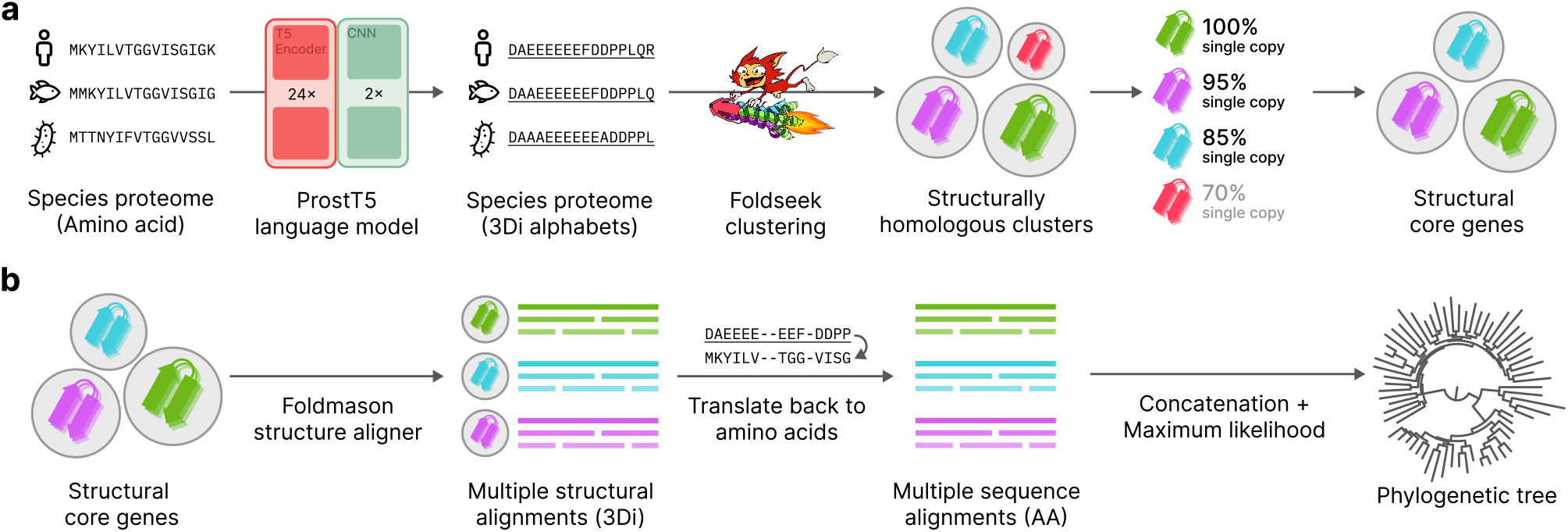
Graphical illustration of the Unicore workflow. (a) The input species proteome, represented as amino acid sequences, is translated to 3Di alphabets (denoted by underscores) using the ProstT5 language model. These 3Di sequences are clustered with Foldseek to group structurally homologous proteins. Structural core genes are identified from these clusters by selecting those conserved as a single-copy in more than a specified proportion of the input species. (b) Foldmason is used to construct multiple structural alignments (MSTAs) for each structural core gene cluster. These alignments are then converted back into amino acid sequences, enabling conventional evolutionary model-based maximum likelihood phylogenetic inference. Finally, a species phylogenetic tree is generated from the concatenated MSAs.

## Methods

Unicore is developed in a modular fashion with Rust programming language. Its modules: `createdb`, `cluster`, `profile`, and `tree` are described in the following sections. For ease of use and automated tree inference, `easy-core` orchestrates these modules into a single workflow.

### Defining structural core genes from species proteomes

Unicore’s first module `createdb` takes a collection of species as an input, where each species is represented as a FASTA file of its proteome (Fig. 1a). These are translated into Foldseek’s 3Di alphabet (12) through the ProstT5 encoder-CNN (26). Both the 3Di sequences and their corresponding amino acid sequences are compiled together into a Foldseek database.

The database is then clustered with the Foldseek `cluster` module, resulting in structural gene clusters. For each cluster, the module `profile` computes the fraction of species out of all input species that have exactly one member structure in that cluster. We denote this fraction as the single-copy coverage of the cluster. Clusters with coverage exceeding a certain threshold (in this study 80% was used as a default value), are defined as the structural core genes.

### Multiple structural alignment and phylogenetic inference

From each structural core gene cluster, the module `tree` converts single-copy members into a Foldmason (29) multiple structural alignment (MSTA; Fig. 1b). Gappy columns, defined as those with a gap fraction of 50% or higher, are then removed from the alignment. Finally, the remaining columns are converted back to amino acid sequences by mapping each 3Di alphabet to its corresponding residue, resulting in multiple sequence alignments (MSAs).

From these structurally-aligned MSAs, Unicore reconstructs the phylogeny by running either IQ-TREE, FastTree, or RAxML (30–32). Unicore produces the maximum likelihood tree inferred from the concatenated MSAs of all identified structural core genes, as well as individual gene trees.

### Tree of life-wide phylogeny using Unicore

We gathered the reference proteome from the UniProt release 2024_02 (33), filtered those with >1,000 proteins and containing >90% of single-copy BUSCOs (v5.4.7) with respective domain used as an OrthoDB v10 lineage (3), resulting in 9,163 proteomes. We then randomly sampled one species from each class, resulting in 166 species comprising all of the Bacterial, Archaeal, and Eukaryotic domains (Supplementary Table 1). We ran Unicore v0-eb516a3 `easy-core` module on these species with default parameters to produce a structural core gene tree with Foldseek v8-ef4e960, Foldmason v1-763a428, and IQ-TREE v2.2.2.3.

### Consistency benchmark - preparation of species sets

Starting with the 9,163 quality-controlled proteomes from the tree of life analysis, we identified 188 taxa ranging from genus to phylum as having at least *N* member species (using *N* = 30 for bacteria and *N* = 20 for fungi), according to NCBI taxonomy annotations (34). We randomly divided each of these 188 taxa into sets of proteomes with exactly *N* members (e.g., fungal taxa with 41 species would yield two sets of 20 and one left out proteome), resulting in 876 bacterial and 98 fungal species sets (Supplementary Table 2). Of these, we randomly sampled one set from each bacterial taxa, resulting in 163 bacterial sets, which were used for the following analyses along with the entirety of 98 fungal sets (Supplementary Table 3).

### Consistency benchmark - comparison with sequence-based methods

We computed structure- and sequence-based core genes from each of the 163 and 98 proteome sets using Unicore `easy-core` and OrthoFinder (v2.5.5), represented as single-copy genes present in at least 90% of the species within each set. This is more stringent than the default threshold, reducing the computational burden in the analysis of sets from lower taxonomic ranks (e.g., genus), where higher fractions of the proteomes are shared among set members. To this end, only the first three modules of Unicore were run with default parameters, except for `profile`, which was run with the option `-t 0.9`. For OrthoFinder, we used the arguments `-og -S mmseqs` and extracted orthogroups that cover at least 90% of species within the dataset.Additionally, BUSCO v5.4.7 with default parameters was used to extract the respective lineage of OrthoDB v10 core genes from each set.

For the core genes defined by each method, we performed HMM-HMM comparisons using `hhsearch` in HH-suite3 v3.3.0 (35) with MAFFT-generated MSAs as input (36). Each MSA was compared against core gene sets from other methods, with a pair classified as shared if the global alignment yielded an *E*-value < 10^*−*3^ with at least 80% coverage. Based on these comparisons, each core gene was classified as shared by both methods or as unique to a specific method.

### Assessment on phylogenetic robustness

We utilized the Unicore, OrthoFinder, and BUSCO core genes of the benchmark species sets from the ‘Consistency benchmark’ section to compare the phylogenetic trees reconstructed from each tool. We used the `tree` module with default parameters to generate phylogenetic trees from the concatenated alignments of Unicore core genes. For the core genes of OrthoFinder and BUSCO, we computed MSAs using MAFFT-linsi v7.525 (36) with default settings, filtered out columns with over 50% gaps, and constructed phylogenetic trees using IQ-TREE v2.3.6 (30) with parameters `-m JTT+F+I+G -B 1000` on the concatenated filtered MSAs. All resulting phylogenetic trees were rooted to yield minimum ancestor deviation (37).

First, we calculated quartet similarity between the trees with tqDist v1.0.2 (38, 39). Then we used the metrics introduced by FoldTree (22), namely taxonomic congruence scores (TCS) and ultrametricity, to further benchmark the quality of the phylogenetic trees.

To test for statistical significance, we performed a *t*-test with the alternative hypothesis that Unicore trees exhibit either higher TCS or lower ultrametricity compared to those generated by OrthoFinder and BUSCO. We controlled for multiple tests by adjusting the *p*-values using the Benjamini-Hochberg method (40) for comparisons with OrthoFinder and BUSCO independently.

### Computation time benchmark

We compared Unicore’s and OrthoFinder’s runtimes on sets of 10, 20, 30, … , 100 species. The species in these sets were randomly sampled from the 330 species of the 11 bacterial sets at the level ‘phyla’ identified as described in the ‘Consistency benchmark’ section (Supplementary Table 3). For Unicore, we used `createdb` with `-max-len 4000 -g` to generate 3Di databases, `cluster` with the arguments `-c “-c 0.8 -max-seqs <maxseqs>“`, where `maxseqs` is set to the larger of 1,000 or 20 times the number of species in the proteome database, and `profile` with default parameters. We separately estimated the time spent on 3Di conversion (`createdb`) and core gene definition (`cluster` + `profile`). Orthofinder was executed with the arguments `-og -S mmseqs`. We used one NVIDIA GeForce RTX 4090 to run Unicore `createdb`, and 128 threads of AMD EPYC 7702P to define core genes using both tools, and took an average runtime of 10 repeated runs. For both tools, we fit a linear regression model between the number of species and the tool’s runtime using R v4.4.1.

## Results and Discussion

### Phylogenetic reconstruction with structurally conserved core genes

We identified 12 single-copy structural core genes with Unicore’s `easy-core` module (Supplementary Table 4), shared by at least 80% out of 166 species spanning the three domains of life (see Methods and Supplementary Table 1). Phylogenetic inference using these structural core genes demonstrated monophyletic delineation of each of the three domains, supported by a bootstrap average of 91.47 (Fig. 2a, see Supplementary Fig. 1 for the detailed tree including species labels). An example of the structural conservation that characterizes Unicore’s core genes is the protein translocase subunit SecY (see Supplementary Table 4 for details), which was not identified as a core gene by OrthoFinder. SecY’s structures are conserved across Archaea, Bacteria, and Eukarya (Fig. 2b) and in each case, predicted 3Di string aligns well with the the experimentally defined structure (TM-score > 0.5).

**Fig. 2.**
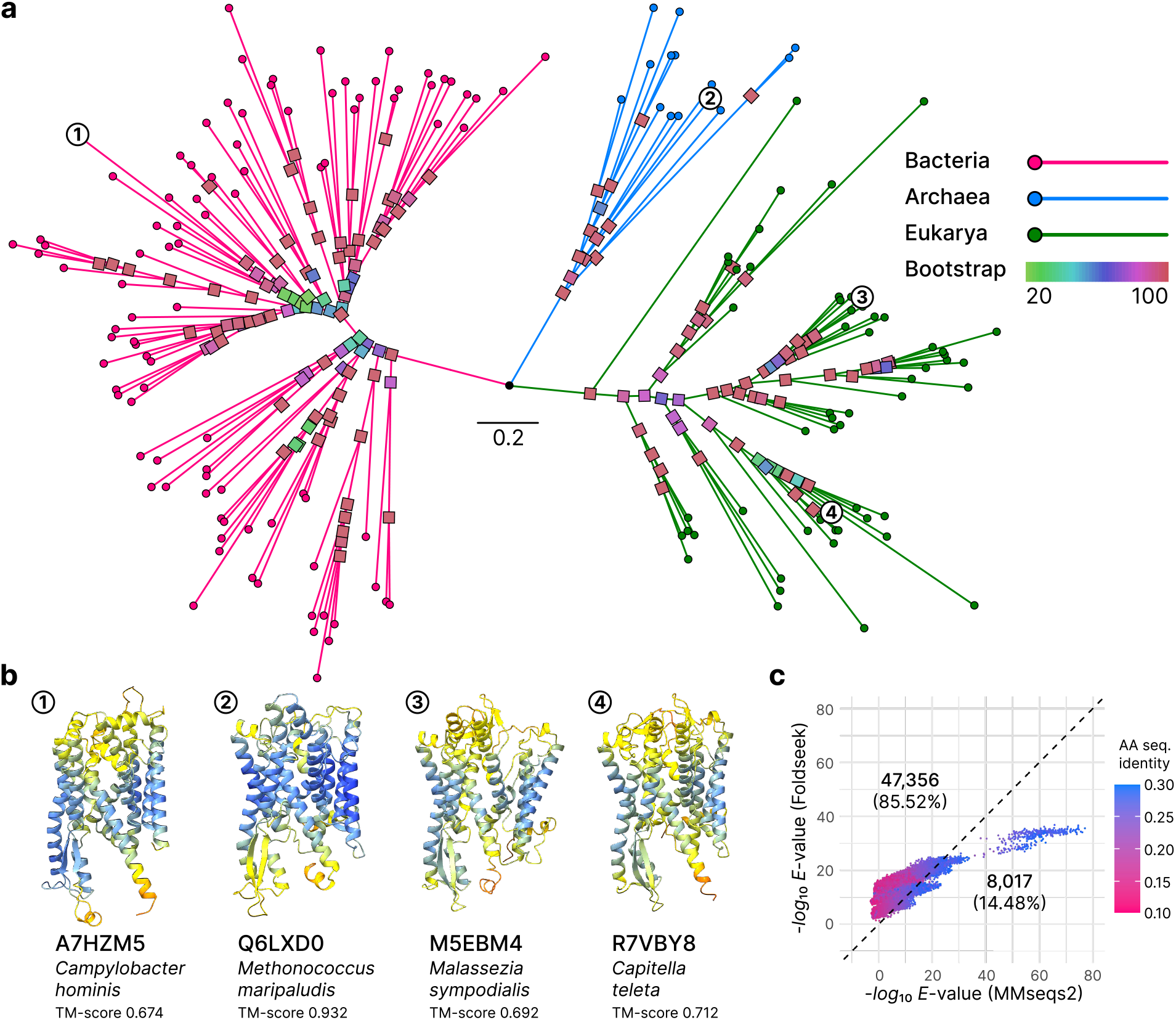
Structural core genes resolve phylogenetic relationships between species with distantly homologous proteins. (a) Phylogenetic tree constructed from 12 structural core genes identified across 166 species, spanning Bacteria (pink), Archaea (blue), and Eukarya (green). Branches are color-coded by domain origin, while internal nodes are colored according to their bootstrap support values. (b) Structural conservation illustrated by SecY protein translocase, one of the identified core genes. Predicted structural models are shown for four species, highlighting structural homology across Bacteria (*Campylobacter hominis*), Archaea (*Methanococcus maripaludis*), Fungi (*Malassezia sympodialis*), and Animalia (*Capitella teleta*), annotated with the TM-score from the alignment against PDB entry *1rhz* chain A. (c) Scatter plot comparing *E* -values derived from MMseqs2 and Foldseek alignments of sequentially distant protein pairs (<30% sequence identity). The data points represent an all-against-all comparison from six structural core genes uniquely defined by Unicore, which were not detected by OrthoFinder in the same species set. The gray dotted line indicates the equivalence *E* (MMseqs2) = *E* (Foldseek), dividing data into two partitions. Counts and proportions for each partition are reported.

Next, we assessed the utility of structural comparison beyond the twilight zone by comparing *E*-values between MM-seqs2’s amino-acid and Foldseek’s 3Di alignments of structural core genes (Fig. 2c). We found that the majority (85.52%) of distantly homologous pairs were better supported (lower *E*-values) by Foldseek, underpinning the capability of 3Di alignments in elucidating distant homology for phylogenetics (20–22).

Here, we applied an 80% threshold for single-copy species coverage to ensure a sufficient number of genes are found to conduct downstream phylogenetic analyses. Adjusting this coverage threshold can impact the resulting set of core genes, which inevitably affects the subsequent phylogenetic analysis (6). Therefore, we made this threshold customizable in Unicore, along with the parameters for Foldseek’s modules, allowing users flexibility in adjusting them according to their specific needs.

### Structural core genes are consistent with sequence-based methods

We evaluated Unicore’s core genes by comparing them with those identified by sequence-based methods, specifically OrthoFinder (9) and BUSCO (41). Out of Unicore’s 170,638 core genes identified in 163 bacterial and 98 fungal sets (see Methods and Supplementary Table 3), 144,658 (84.75%) were detected as such by both OrthoFinder and BUSCO (Supplementary Fig. 2a), while 3,142 (1.84%) genes were unique to Unicore. In the opposite direction, we found 90.53% of OrthoFinder’s genes and 91.50% of BUSCO’s genes were detectable by Unicore (Supplementary Fig. 2b and 2c), further supporting the validity of Unicore’s core genes.

### Robustness of the phylogenetic trees reconstructed with Unicore

We further assessed the quality of phylogenetic inference with Unicore structural core genes identified in the 163 bacterial and 98 fungal benchmark sets. Unicore’s bacterial trees showed median quartet similarity to OrthoFinder’s and BUSCO’s trees was 0.985 and 0.968, respectively (Fig. 3a, left). This measure was even higher in the fungal dataset, reaching 1.0 compared to both methods (Fig. 3a, right). However, this may result from having smaller fungal sets than bacterial sets (see Methods), an artifact due to the limited availability of high-quality fungal genomes.

**Fig. 3.**
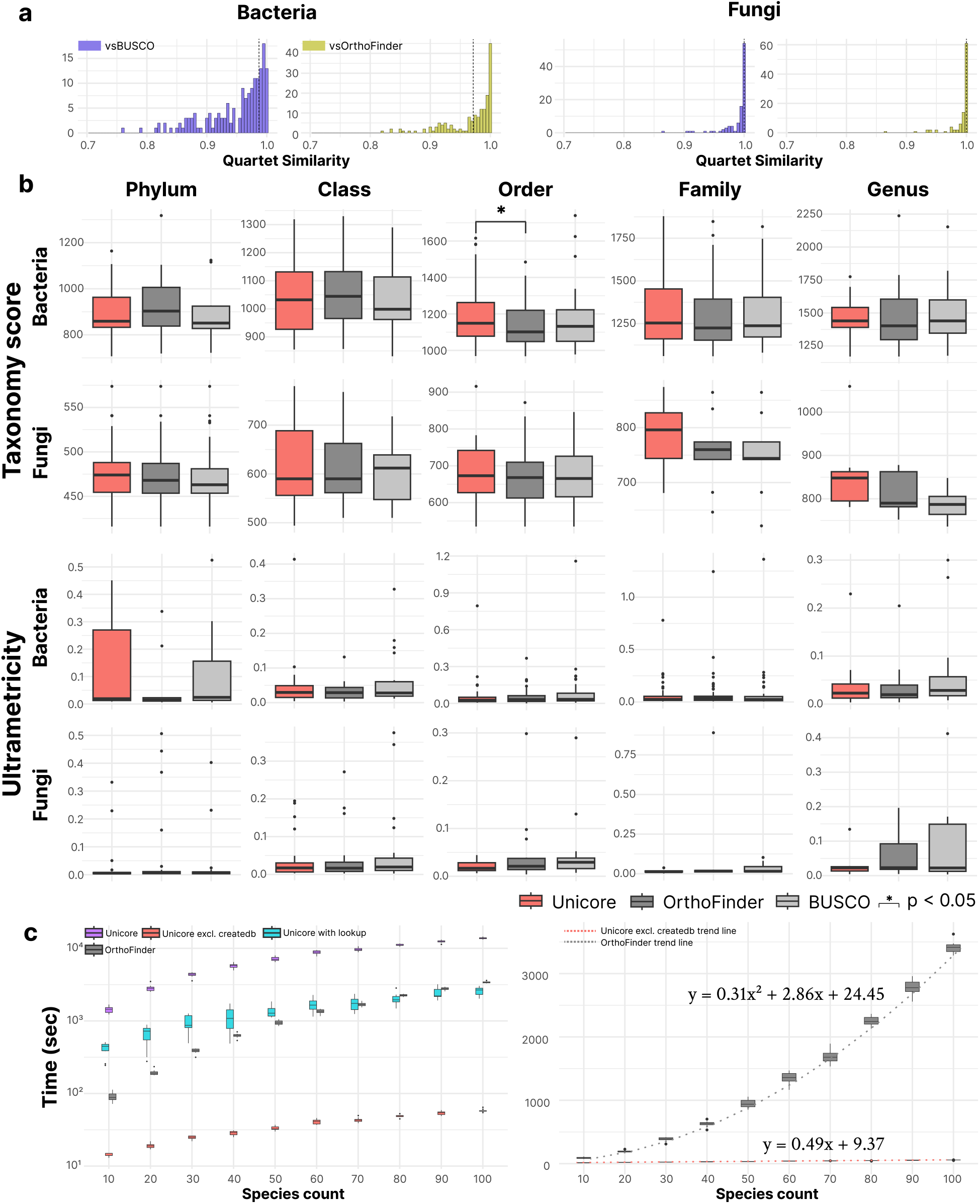
Benchmark results comparing the speed and phylogenetic robustness of trees generated by Unicore, OrthoFinder, and BUSCO. (a) The histogram shows quartet similarity scores for trees computed with Unicore against OrthoFinder (yellow) and BUSCO (blue). The left panel displays bacterial trees, while the right shows fungal trees. A bin width of 0.005 is used, with the median marked by a vertical dotted line. (b) TCS and ultrametricity for Unicore (red), OrthoFinder (dark gray), and BUSCO (gray) are compared across taxonomic ranks. The Benjamini-Hochberg method corrected p-values, with significant results (p < 0.05) indicated by a star. (c) Left: Unicore (purple) includes the entire process from predicting 3Di sequences to core gene inference. Unicore with lookup (cyan) accounts only for predicting 3Di sequences absent in the AlphaFold database, while Unicore excl. createdb (red) benchmarks core gene inference alone. OrthoFinder (dark gray) measures orthogroup inference. Right: Results are plotted on a linear y-axis. Core gene inference with Unicore exhibits linear time complexity, unlike OrthoFinder’s quadratic scaling, as shown by the dotted trend line. Regression analysis confirmed this, with Unicore fitting poorly to a quadratic model (coefficient = -0.0002, *p* = 0.572) but strongly to a linear model (coefficient = 0.4932, *p* < 10^*−*15^). In contrast, OrthoFinder’s runtime fits well with a quadratic model (coefficient = 0.3095, *p* < 10^*−*15^).

Meanwhile, Unicore trees showed comparable taxonomic congruence and ultrametricity with the trees computed based on the core genes of OrthoFinder and BUSCO, across all taxonomic ranks for both bacteria and fungi (Fig. 3b). Put together, structural core genes of Unicore reconstructed robust species phylogeny comparable to the sequence-only methods, highlighting both the validity of the structure-based phylogenetic approach and its prospect of elucidating beyond the twilight zone of amino acid sequences.

### Unicore allows structure-based phylogeny at scale

To demonstrate the scalability of our method, we compared the time it takes Unicore and OrthoFinder to detect core genes in benchmark sets of species. Excluding database creation, Unicore scales linearly with the number of input species, thereby being faster than the quadratically-scaling OrthoFinder, reaching a factor of 58.69 in runtime over 100 species (Fig. 3c). However, database creation is Unicore’s biggest bottleneck, which requires 3Di conversion by ProstT5, making it up to 4.05 times slower than OrthoFinder. To alleviate this, we implemented a sequence lookup method that rapidly retrieves 3Di sequences from the AFDB (27) for exact sequence matches, thereby improving Unicore’s runtime by up to 5.44-fold, which was 1.34-fold faster than Orthofinder.

Since conversion with ProstT5 heavily depends on the model and amount of GPU resources, the overall throughput is expected to be improved concurrent to the rapid development of graphics hardware. In this context, owing to its quasi-linear clustering algorithm (16), Unicore’s scalability will become increasingly powerful against the ever-increasing flood of biological data.

### Concluding remarks

Unicore implements a large-scale, structure-based phylogenetic method powered by state-of-the-art tools for protein structural analysis, compiled into a fully automated bioinformatics pipeline. Accompanied by the recent developments in structural phylogenetics (20–22, 29, 42), we anticipate our tool to further expand the utility of protein structures from an evolutionary perspective.

## Data Availability

Unicore is implemented in Rust and is available as GPLv3 licensed free open-source software at https://github.com/steineggerlab/unicore.

## Acknowledgments

M.S. acknowledges support by the National Research Foundation of Korea grants (2020M3-A9G7-103933, 2021-R1C1-C102065 and 2021-M3A9-I4021220, RS-2024-00396026), Samsung DS research fund, Creative-Pioneering Researchers Program and AI-Bio Research Grant through Seoul National University. We thank Eli Levy Karin of ELKMO, as well as Rachel Seongeun Kim, Hyunbin Kim, and Jaebeom Kim of the Steinegger Lab for their help with revising the manuscript.

