## Supplementary material for "Unicore Enables Scalable and Accurate Phylogenetic Reconstruction with Structural Core Genes"

[1]Interdisciplinary Program in Bioinformatics, Seoul National University, Seoul 08826, Republic of Korea, [2]School of Biological Sciences, Seoul National University, Seoul 08826, Republic of Korea, [3]Institute of Molecular Biology and Genetics, Seoul National University, Seoul 08826, Republic of Korea, [4]Artificial Intelligence Institute, Seoul National University, Seoul 08826, Republic of Korea

### SUPPLEMENTARY MATERIALS

#### Figures

- Supplementary Fig. 1. Structural core gene tree of 166 species across the tree of life.
- Supplementary Fig. 2. Venn diagram representation of the overlap between the orthologs.

#### Tables

- Supplementary Table 1. Taxonomic description of 166 species spanning across the tree of life.
- Supplementary Table 2. List of the 188 taxa and 974 sub-sampled sets of species defined from 9,163 quality-controlled proteomes.
- Supplementary Table 3. Detailed list of the 261 sets on which benchmark was performed.
- Supplementary Table 4. List of 12 structural core genes defined from 166 species spanning the tree of life.

†These authors contributed equally.

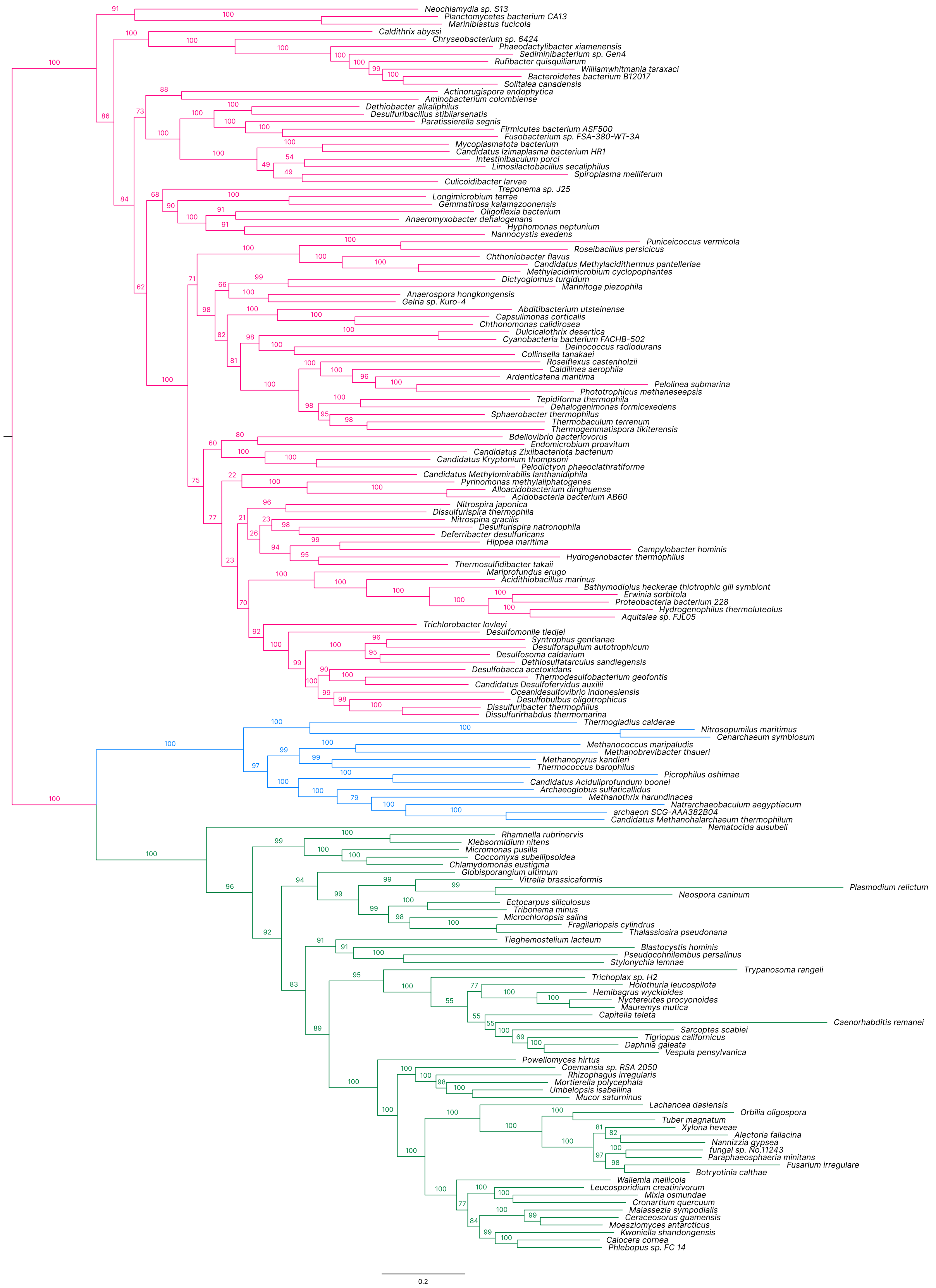


**Supplementary Fig. 1.** Structural core gene tree of 166 species across the tree of life. Branches and bootstrap support values are colored by their originating domain (magenta, Bacteria; cyan, Archaea; green, Eukarya).

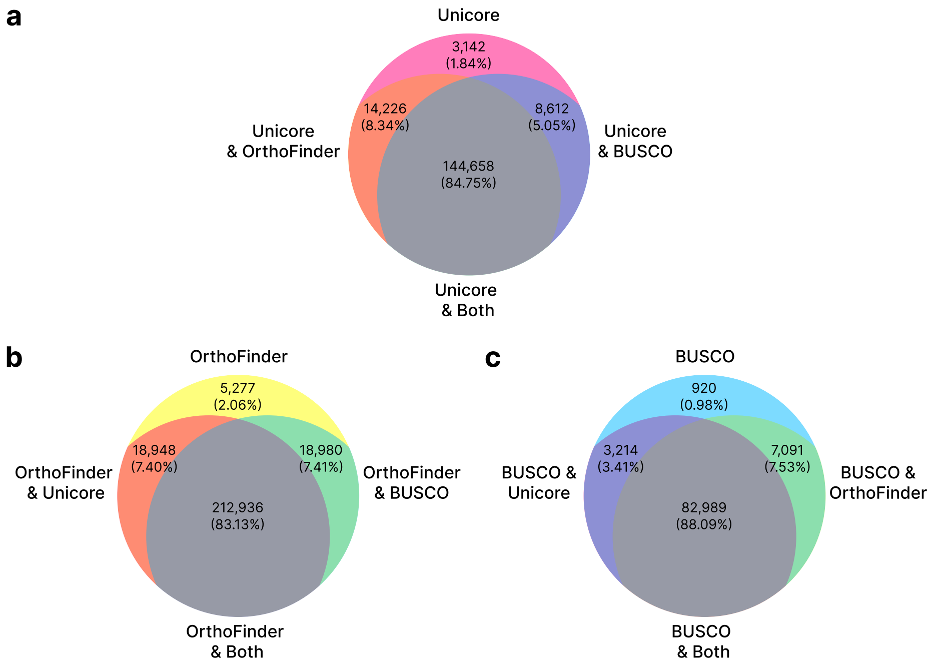


**Supplementary Fig. 2.** Venn diagram representation of the overlap between the orthologs. Ratios of the orthologs shared with none, either or both of the other tools to the total number of orthologs are shown for (a) Unicore, (b) OrthoFinder and (c) BUSCO.

**Supplementary Table 1.** Taxonomic description of 166 species spanning across the tree of life. For each species we described its class, from which it was randomly selected among the members, along with its UniProt identifier of reference proteome.

| Identifier | Species name | Class name |
|---|---|---|
| UP000000263 | *Roseiflexus castenholzii* | Chloroflexia |
| UP000000323 | *Thermobaculum terrenum* | Chloroflexota |
| UP000000442 | *Desulforapulum autotrophicum* | Desulfobacteria |
| UP000000483 | *Desulfobacca acetoxidans* | Desulfobaccia |
| UP000000590 | *Methanococcus maripaludis* | Methanococci |
| UP000000758 | *Cenarchaeum symbiosum* | Nitrososphaerota |
| UP000000792 | *Nitrosopumilus maritimus* | Nitrososphaeria |
| UP000001400 | *Aciduliprofundum boonei** | Thermoplasmatota* |
| UP000001449 | *Thalassiosira pseudonana* | Coscinodiscophyceae |
| UP000001520 | *Deferribacter desulfuricans* | Deferribacteres |
| UP000001826 | *Methanopyrus kandleri* | Methanopyri |
| UP000001876 | *Micromonas pusilla* | Mamiellophyceae |
| UP000001935 | *Anaeromyxobacter dehalogenans* | Myxococcia |
| UP000001959 | *Hyphomonas neptunium* | Alphaproteobacteria |
| UP000002027 | *Sphaerobacter thermophilus* | Thermomicrobia |
| UP000002366 | *Aminobacterium colombiense* | Synergistia |
| UP000002407 | *Campylobacter hominis* | Epsilonproteobacteria |
| UP000002420 | *Trichlorobacter lovleyi* | Desulfuromonadia |
| UP000002524 | *Deinococcus radiodurans* | Deinococci |
| UP000002574 | *Hydrogenobacter thermophilus* | Aquificae |
| UP000002630 | *Ectocarpus siliculosus* | Phaeophyceae |
| UP000002669 | *Nannizzia gypsea* | Eurotiomycetes |
| UP000002724 | *Pelodictyon phaeoclathratiforme* | Chlorobiia |
| UP000004671 | *Caldithrix abyssi* | Calditrichia |
| UP000004830 | *Collinsella tanakaei* | Coriobacteriia |
| UP000005242 | *Wallemia mellicola* | Wallemiomycetes |
| UP000005270 | *Thermogladius calderae* | Thermoprotei |
| UP000005824 | *Chthoniobacter flavus* | Spartobacteria |
| UP000005877 | *Methanothrix harundinacea* | Methanomicrobia |
| UP000006055 | *Desulfomonile tiedjei* | Desulfomonilia |
| UP000006443 | *Dethiobacter alkaliphilus* | Dethiobacteria |
| UP000006583 | *Thermodesulfobacterium geofontis* | Thermodesulfobacteria |
| UP000007161 | *Marinitoga piezophila* | Thermotogae |
| UP000007264 | *Coccomyxa subellipsoidea* | Trebouxiophyceae |
| UP000007478 | *Thermococcus barophilus* | Thermococci |
| UP000007494 | *Neospora caninum* | Conoidasida |
| UP000007590 | *Solitalea canadensis* | Sphingobacteriia |
| UP000007719 | *Dictyoglomus turgidum* | Dictyoglomia |
| UP000007880 | *Caldilinea aerophila* | Caldilineae |
| UP000008139 | *Hippea maritima* | Desulfurellia |
| UP000008281 | *Caenorhabditis remanei* | Chromadorea |
| UP000008784 | *Orbilia oligospora* | Orbiliomycetes |
| UP000009131 | *Mixia osmundae* | Mixiomycetes |
| UP000011704 | *Nitrospina gracilis* | Nitrospinia |
| UP000011976 | *Moesziomyces antarcticus* | Ustilaginomycetes |
| UP000013307 | *Archaeoglobus sulfaticallidus* | Archaeoglobi |
| UP000014227 | *Chthonomonas calidirosea* | Chthonomonadetes |
| UP000014760 | *Capitella teleta* | Polychaeta |
| UP000017395 | *Firmicutes* bacterium ASF500* | Bacillota |
| UP000019132 | *Globisporangium ultimum* | Oomycota |
| UP000019151 | *Gemmatirosa kalamazoonensis* | Gemmatimonadetes |
| UP000029408 | *Izimaplasma* bacterium HR1* | Izimaplasma* |
| UP000029736 | *Phaeodactylibacter xiamenensis* | Saprospiria |
| UP000031518 | *Pyrinomonas methylaliphatogenes* | Blastocatellia |
| UP000031737 | *Trypanosoma rangeli* | Kinetoplastea |
| UP000032233 | *Dethiosulfatarculus sandiegensis* | Desulfarculia |
| UP000035337 | *Endomicrobium proavitum* | Endomicrobiia |
| UP000037784 | *Ardenticatena maritima* | Ardenticatenia |
| UP000039865 | *Stylonychia lemnae* | Spirotrichea |
| UP000041254 | *Vitrella brassicaformis* | Eukaryota |
| UP000050934 | *Limosilactobacillus secaliphilus* | Bacilli |
| UP000054361 | fungal sp. No.11243* | Unidentified Fungi* |

**Supplementary Table 1.** (Continued.)

| Identifier | Species name | Class name |
|---|---|---|
| UP000054524 | *Nematocida ausubeli* | Microsporidia |
| UP000054558 | *Klebsormidium nitens* | Klebsormidiophyceae |
| UP000054937 | *Pseudocohnilembus persalinus* | Oligohymenophorea |
| UP000063234 | *Thermosulfidibacter takaii* | Thermosulfidibacteria |
| UP000070412 | *Sarcoptes scabiei* | Arachnida |
| UP000070560 | *Desulfofervidus auxilii** | Desulfofervidia* |
| UP000075320 | *Bdellovibrio bacteriovorus* | Bdellovibrionia |
| UP000076078 | *Tieghemostelium lacteum* | Eumycetozoa |
| UP000076632 | *Xylona heveae* | Xylonomycetes |
| UP000076842 | *Calocera cornea* | Dacrymycetes |
| UP000078348 | *Blastocystis hominis* | Bigyra |
| UP000093080 | *Dissulfuribacter thermophilus* | Dissulfuribacteria |
| UP000095255 | *Desulfuribacillus stibiiarsenatis* | Desulfuribacillia |
| UP000095751 | *Fragilariopsis cylindrus* | Bacillariophyceae |
| UP000182011 | *Kryptonium thompsoni** | Kryptonia* |
| UP000185744 | *Methanohalarchaeum thermophilum** | Methanonatronarchaeia |
| UP000185934 | *Dehalogenimonas formicexedens* | Dehalococcoidia |
| UP000186303 | *Malassezia sympodialis* | Malasseziomycetes |
| UP000190274 | *Lachancea dasiensis* | Saccharomycetes |
| UP000192042 | *Nitrospira japonica* | Nitrospiria |
| UP000192315 | *Picrophilus oshimae* | Thermoplasmata |
| UP000193467 | *Leucosporidium creatinivorum* | Microbotryomycetes |
| UP000198744 | *Syntrophus gentianae* | Syntrophia |
| UP000199400 | *Nannocystis exedens* | Myxococcota |
| UP000199452 | *Williamwhitmania taraxaci* | Bacteroidia |
| UP000216329 | *Bacteroidetes* bacterium B1* | Bacteroidota |
| UP000220158 | *Plasmodium relictum* | Aconoidasida |
| UP000223071 | *Tepidiforma thermophila* | Tepidiformia |
| UP000232323 | *Chlamydomonas eustigma* | Chlorophyceae |
| UP000234329 | *Acidithiobacillus marinus* | Acidithiobacillia |
| UP000237684 | *Abditibacterium utsteinense* | Abditibacteriia |
| UP000238196 | *Proteobacteria* bacterium 228* | Pseudomonadota |
| UP000240531 | Archaeon SCG-AAA382B04* | Unidentified Archaea* |
| UP000245783 | *Ceraceosorus guamensis* | Exobasidiomycetes |
| UP000246991 | *Tuber magnatum* | Pezizomycetes |
| UP000248706 | *Thermogemmatispora tikiterensis* | Ktedonobacteria |
| UP000250088 | *Natrarchaeobaculum aegyptiacum* | Halobacteria |
| UP000251717 | *Methanobrevibacter thaueri* | Methanobacteria |
| UP000253843 | *Trichoplax* sp. H2* | Uniplacotomia |
| UP000256388 | *Pelolinea submarina* | Anaerolineae |
| UP000262004 | *Hydrogenophilus thermoluteolus* | Hydrogenophilia |
| UP000268059 | *Intestinibaculum porci* | Erysipelotrichia |
| UP000270538 | *Chryseobacterium* sp. 6424* | Flavobacteriia |
| UP000271624 | *Dulcicalothrix desertica* | Cyanophyceae |
| UP000272943 | *Aquitalea* sp. FJL05* | Betaproteobacteria |
| UP000276223 | *Desulfosoma caldarium* | Syntrophobacteria |
| UP000287394 | *Capsulimonas corticalis* | Armatimonadia |
| UP000289195 | *Neochlamydia* sp. S13* | Chlamydiia |
| UP000294625 | *Treponema* sp. J25* | Spirochaetia |
| UP000295063 | *Anaerospora hongkongensis* | Negativicutes |
| UP000295281 | *Actinorugispora endophytica* | Actinomycetes |
| UP000297299 | *Botryotinia calthae* | Leotiomycetes |
| UP000298715 | *Spiroplasma melliferum* | Mollicutes |
| UP000306585 | *Mariprofundus erugo* | Zetaproteobacteria |
| UP000306912 | *Culicoidibacter larvae* | Culicoidibacteria |
| UP000309626 | *Bathymodiolus heckerae* thiotrophic gill symbiont* | Unidentified Bacteria* |
| UP000315010 | *Planctomycetes* bacterium CA13* | Planctomycetota |
| UP000318571 | *Tigriopus californicus* | Hexanauplia |
| UP000318582 | *Powellomyces hirtus* | Chytridiomycetes |
| UP000322214 | *Mariniblastus fucicola* | Planctomycetia |
| UP000322225 | *Kwoniella shandongensis* | Tremellomycetes |
| UP000324789 | *Acidobacteria* bacterium AB60* | Acidobacteriota |
| UP000334340 | *Methylomirabilis lanthanidiphila** | Division NC10* |
| UP000355283 | *Microchloropsis salina* | Eustigmatophyceae |

**Supplementary Table 1.** (Continued.)

| Identifier | Species name | Class name |
|---|---|---|
| UP000381693 | *Methylacidimicrobium cyclopophantes* | Verrucomicrobiota |
| UP000412907 | *Zixiibacteriota* bacterium* | Zixiibacteriota* |
| UP000424752 | *Erwinia sorbitola* | Gammaproteobacteria |
| UP000448292 | *Oceanidesulfovibrio indonesiensis* | Desulfovibrionia |
| UP000469346 | *Dissulfurirhabdus thermomarina* | Deltaproteobacteria |
| UP000515312 | *Alloacidobacterium dinghuense* | Terriglobia |
| UP000516360 | *Dissulfurispira thermophila* | Thermodesulfovibrionia |
| UP000525652 | *Puniceicoccus vermicola* | Opitutae |
| UP000528322 | *Desulfurispira natronophila* | Chrysiogenetes |
| UP000563094 | *Rufibacter quisquiliarum* | Cytophagia |
| UP000579846 | *Sediminibacterium* sp. Gen4* | Chitinophagia |
| UP000582837 | *Longimicrobium terrae* | Longimicrobiia |
| UP000591197 | *Fusobacterium* sp. FSA-380-WT-3A* | Fusobacteriia |
| UP000594468 | *Phototrophicus methaneseepsis* | Thermofonsia* |
| UP000596092 | *Desulfobulbus oligotrophicus* | Desulfobulbia |
| UP000600918 | *Vespula pensylvanica* | Insecta |
| UP000601171 | *Paratissierella segnis* | Tissierellia |
| UP000603453 | *Mucor saturninus* | Mucoromycetes |
| UP000610699 | *Oligoflexia bacterium* | Oligoflexia |
| UP000627505 | *Cyanobacteria* bacterium FACHB-502* | Cyanobacteriota |
| UP000644507 | *Roseibacillus persicicus* | Verrucomicrobiae |
| UP000645828 | *Nyctereutes procyonoides* | Mammalia |
| UP000654370 | *Umbelopsis isabellina* | Umbelopsidomycetes |
| UP000663859 | *Methylacidithermus pantelleriae** | Methylacidiphilae |
| UP000664203 | *Alectoria fallacina* | Lecanoromycetes |
| UP000664859 | *Tribonema minus* | Xanthophyceae |
| UP000678917 | *Mycoplasmatota bacterium* | Mycoplasmatota |
| UP000726737 | *Mortierella polycephala* | Mortierellomycetes |
| UP000756921 | *Paraphaeosphaeria minitans* | Dothideomycetes |
| UP000789390 | *Daphnia galeata* | Branchiopoda |
| UP000796880 | *Rhamnella rubrinervis* | Magnoliopsida |
| UP000813600 | *Phlebopus* sp. FC_14* | Agaricomycetes |
| UP000824219 | *Hemibagrus wyckioides* | Actinopteri |
| UP000826003 | *Gelria* sp. Kuro-4* | Clostridia |
| UP000827986 | *Mauremys mutica* | Chordata |
| UP000886653 | *Cronartium quercuum* | Pucciniomycetes |
| UP001151568 | *Coemansia* sp. RSA 2050* | Kickxellomycetes |
| UP001152130 | *Fusarium irregulare* | Sordariomycetes |
| UP000018888 | *Rhizophagus irregularis* | Glomeromycetes |
| UP001152320 | *Holothuria leucospilota* | Holothuroidea |

*Candidate or unidentified classification

**Supplementary Table 2.** List of the 188 taxa and 974 sub-sampled sets of species defined from 9,163 quality-controlled proteomes. We provided corresponding NCBI taxonomy ID, rank and type (Bacteria or Fungi), along with the number of member species and sub-sampled sets.

| ID | Scientific name | Rank | Type | Members | Sets |
|---|---|---|---|---|---|
| 136 | Spirochaetales | Order | Bacteria | 35 | 1 |
| 237 | *Flavobacterium* | Genus | Bacteria | 73 | 2 |
| 265 | *Paracoccus* | Genus | Bacteria | 36 | 1 |
| 286 | *Pseudomonas* | Genus | Bacteria | 55 | 1 |
| 356 | Hyphomicrobiales | Order | Bacteria | 401 | 13 |
| 379 | *Rhizobium* | Genus | Bacteria | 34 | 1 |
| 433 | Acetobacteraceae | Family | Bacteria | 73 | 2 |
| 468 | Moraxellaceae | Family | Bacteria | 34 | 1 |
| 481 | Neisseriaceae | Family | Bacteria | 35 | 1 |
| 506 | Alcaligenaceae | Family | Bacteria | 50 | 1 |
| 543 | Enterobacteriaceae | Family | Bacteria | 41 | 1 |
| 641 | Vibrionaceae | Family | Bacteria | 48 | 1 |
| 662 | *Vibrio* | Genus | Bacteria | 35 | 1 |
| 815 | Bacteroidaceae | Family | Bacteria | 33 | 1 |
| 838 | *Prevotella* | Genus | Bacteria | 40 | 1 |

**Supplementary Table 2.** (Continued.)

| ID | Scientific name | Rank | Type | Members | Sets |
|---|---|---|---|---|---|
| 976 | Bacteroidota | Phylum | Bacteria | 894 | 29 |
| 1117 | Cyanobacteriota | Phylum | Bacteria | 178 | 5 |
| 1161 | Nostocales | Order | Bacteria | 44 | 1 |
| 1224 | Pseudomonadota | Phylum | Bacteria | 2733 | 91 |
| 1236 | Gammaproteobacteria | Class | Bacteria | 886 | 29 |
| 1239 | Bacillota | Phylum | Bacteria | 1493 | 49 |
| 1268 | Micrococcaceae | Family | Bacteria | 94 | 3 |
| 1300 | Streptococcaceae | Family | Bacteria | 52 | 1 |
| 1301 | *Streptococcus* | Genus | Bacteria | 40 | 1 |
| 1385 | Bacillales | Order | Bacteria | 559 | 18 |
| 1386 | *Bacillus* | Genus | Bacteria | 62 | 2 |
| 1485 | *Clostridium* | Genus | Bacteria | 105 | 3 |
| 1653 | Corynebacteriaceae | Family | Bacteria | 74 | 2 |
| 1663 | *Arthrobacter* | Genus | Bacteria | 40 | 1 |
| 1678 | *Bifidobacterium* | Genus | Bacteria | 43 | 1 |
| 1716 | *Corynebacterium* | Genus | Bacteria | 74 | 2 |
| 1760 | Actinomycetes | Class | Bacteria | 1762 | 58 |
| 1762 | Mycobacteriaceae | Family | Bacteria | 75 | 2 |
| 1763 | *Mycobacterium* | Genus | Bacteria | 40 | 1 |
| 1817 | *Nocardia* | Genus | Bacteria | 34 | 1 |
| 1839 | *Nocardioides* | Genus | Bacteria | 73 | 2 |
| 1873 | *Micromonospora* | Genus | Bacteria | 41 | 1 |
| 1883 | *Streptomyces* | Genus | Bacteria | 353 | 11 |
| 2004 | Streptosporangiaceae | Family | Bacteria | 59 | 1 |
| 2012 | Thermomonosporaceae | Family | Bacteria | 33 | 1 |
| 2037 | Actinomycetales | Order | Bacteria | 44 | 1 |
| 2049 | Actinomycetaceae | Family | Bacteria | 44 | 1 |
| 2062 | Streptomycetaceae | Family | Bacteria | 380 | 12 |
| 2070 | Pseudonocardiaceae | Family | Bacteria | 114 | 3 |
| 4890 | Ascomycota | Phylum | Fungi | 527 | 26 |
| 4891 | Saccharomycetes | Class | Fungi | 71 | 3 |
| 4892 | Saccharomycetales | Order | Fungi | 71 | 3 |
| 4893 | Saccharomycetaceae | Family | Fungi | 23 | 1 |
| 5042 | Eurotiales | Order | Fungi | 111 | 5 |
| 5052 | *Aspergillus* | Genus | Fungi | 59 | 2 |
| 5073 | *Penicillium* | Genus | Fungi | 42 | 2 |
| 5125 | Hypocreales | Order | Fungi | 104 | 5 |
| 5178 | Helotiales | Order | Fungi | 28 | 1 |
| 5204 | Basidiomycota | Phylum | Fungi | 153 | 7 |
| 5338 | Agaricales | Order | Fungi | 47 | 2 |
| 5455 | *Colletotrichum* | Genus | Fungi | 22 | 1 |
| 5506 | *Fusarium* | Genus | Fungi | 51 | 2 |
| 13687 | *Sphingomonas* | Genus | Bacteria | 85 | 2 |
| 28056 | Micromonosporaceae | Family | Bacteria | 104 | 3 |
| 28211 | Alphaproteobacteria | Class | Bacteria | 1302 | 43 |
| 28216 | Betaproteobacteria | Class | Bacteria | 534 | 17 |
| 28256 | Halomonadaceae | Family | Bacteria | 37 | 1 |
| 29547 | Campylobacterota | Phylum | Bacteria | 82 | 2 |
| 31953 | Bifidobacteriaceae | Family | Bacteria | 48 | 1 |
| 31957 | Propionibacteriaceae | Family | Bacteria | 40 | 1 |
| 31979 | Clostridiaceae | Family | Bacteria | 130 | 4 |
| 31989 | Paracoccaceae | Family | Bacteria | 200 | 6 |
| 32003 | Nitrosomonadales | Order | Bacteria | 61 | 2 |
| 32033 | Xanthomonadaceae | Family | Bacteria | 76 | 2 |
| 33882 | *Microbacterium* | Genus | Bacteria | 68 | 2 |
| 33958 | Lactobacillaceae | Family | Bacteria | 124 | 4 |
| 41297 | Sphingomonadaceae | Family | Bacteria | 191 | 6 |
| 44249 | *Paenibacillus* | Genus | Bacteria | 118 | 3 |
| 49546 | Flavobacteriaceae | Family | Bacteria | 320 | 10 |
| 57723 | Acidobacteriota | Phylum | Bacteria | 32 | 1 |
| 69277 | Phyllobacteriaceae | Family | Bacteria | 64 | 2 |
| 72274 | Pseudomonadales | Order | Bacteria | 85 | 2 |
| 72275 | Alteromonadaceae | Family | Bacteria | 34 | 1 |
| 72293 | Helicobacteraceae | Family | Bacteria | 30 | 1 |

**Supplementary Table 2.** (Continued.)

| ID | Scientific name | Rank | Type | Members | Sets |
|---|---|---|---|---|---|
| 74201 | Verrucomicrobiota | Phylum | Bacteria | 43 | 1 |
| 75682 | Oxalobacteraceae | Family | Bacteria | 66 | 2 |
| 76892 | Caulobacteraceae | Family | Bacteria | 36 | 1 |
| 80840 | Burkholderiales | Order | Bacteria | 372 | 12 |
| 80864 | Comamonadaceae | Family | Bacteria | 128 | 4 |
| 81852 | Enterococcaceae | Family | Bacteria | 46 | 1 |
| 82115 | Rhizobiaceae | Family | Bacteria | 67 | 2 |
| 84566 | Sphingobacteriaceae | Family | Bacteria | 110 | 3 |
| 84567 | *Pedobacter* | Genus | Bacteria | 34 | 1 |
| 84998 | Coriobacteriia | Class | Bacteria | 47 | 1 |
| 85004 | Bifidobacteriales | Order | Bacteria | 48 | 1 |
| 85006 | Micrococcales | Order | Bacteria | 474 | 15 |
| 85007 | Mycobacteriales | Order | Bacteria | 243 | 8 |
| 85008 | Micromonosporales | Order | Bacteria | 104 | 3 |
| 85009 | Propionibacteriales | Order | Bacteria | 147 | 4 |
| 85010 | Pseudonocardiales | Order | Bacteria | 114 | 3 |
| 85011 | Kitasatosporales | Order | Bacteria | 380 | 12 |
| 85012 | Streptosporangiales | Order | Bacteria | 115 | 3 |
| 85015 | Nocardioidaceae | Family | Bacteria | 96 | 3 |
| 85016 | Cellulomonadaceae | Family | Bacteria | 33 | 1 |
| 85021 | Intrasporangiaceae | Family | Bacteria | 33 | 1 |
| 85023 | Microbacteriaceae | Family | Bacteria | 220 | 7 |
| 85025 | Nocardiaceae | Family | Bacteria | 67 | 2 |
| 85030 | Geodermatophilaceae | Family | Bacteria | 31 | 1 |
| 91061 | Bacilli | Class | Bacteria | 825 | 27 |
| 91347 | Enterobacterales | Order | Bacteria | 106 | 3 |
| 92860 | Pleosporales | Order | Fungi | 46 | 2 |
| 110618 | Nectriaceae | Family | Fungi | 57 | 2 |
| 117743 | Flavobacteriia | Class | Bacteria | 373 | 12 |
| 117747 | Sphingobacteriia | Class | Bacteria | 110 | 3 |
| 119045 | Methylobacteriaceae | Family | Bacteria | 34 | 1 |
| 119060 | Burkholderiaceae | Family | Bacteria | 89 | 2 |
| 135613 | Chromatiales | Order | Bacteria | 71 | 2 |
| 135614 | Xanthomonadales | Order | Bacteria | 114 | 3 |
| 135619 | Oceanospirillales | Order | Bacteria | 110 | 3 |
| 135620 | Oceanospirillaceae | Family | Bacteria | 44 | 1 |
| 135621 | Pseudomonadaceae | Family | Bacteria | 71 | 2 |
| 135622 | Alteromonadales | Order | Bacteria | 118 | 3 |
| 135623 | Vibrionales | Order | Bacteria | 48 | 1 |
| 147541 | Dothideomycetes | Class | Fungi | 98 | 4 |
| 147545 | Eurotiomycetes | Class | Fungi | 143 | 7 |
| 147548 | Leotiomycetes | Class | Fungi | 32 | 1 |
| 147550 | Sordariomycetes | Class | Fungi | 164 | 8 |
| 155619 | Agaricomycetes | Class | Fungi | 97 | 4 |
| 165695 | *Sphingobium* | Genus | Bacteria | 30 | 1 |
| 165696 | *Novosphingobium* | Genus | Bacteria | 40 | 1 |
| 171549 | Bacteroidales | Order | Bacteria | 166 | 5 |
| 171552 | Prevotellaceae | Family | Bacteria | 64 | 2 |
| 186801 | Clostridia | Class | Bacteria | 557 | 18 |
| 186802 | Eubacteriales | Order | Bacteria | 321 | 10 |
| 186803 | Lachnospiraceae | Family | Bacteria | 131 | 4 |
| 186806 | Eubacteriaceae | Family | Bacteria | 33 | 1 |
| 186817 | Bacillaceae | Family | Bacteria | 293 | 9 |
| 186818 | Planococcaceae | Family | Bacteria | 42 | 1 |
| 186822 | Paenibacillaceae | Family | Bacteria | 159 | 5 |
| 186826 | Lactobacillales | Order | Bacteria | 266 | 8 |
| 194924 | Desulfovibrionaceae | Family | Bacteria | 32 | 1 |
| 200643 | Bacteroidia | Class | Bacteria | 189 | 6 |
| 200644 | Flavobacteriales | Order | Bacteria | 373 | 12 |
| 200666 | Sphingobacteriales | Order | Bacteria | 110 | 3 |
| 200940 | Thermodesulfobacteriota | Phylum | Bacteria | 117 | 3 |
| 201174 | Actinomycetota | Phylum | Bacteria | 1809 | 60 |
| 203682 | Planctomycetota | Phylum | Bacteria | 73 | 2 |
| 203683 | Planctomycetia | Class | Bacteria | 73 | 2 |

**Supplementary Table 2.** (Continued.)

| ID | Scientific name | Rank | Type | Members | Sets |
|---|---|---|---|---|---|
| 203691 | Spirochaetota | Phylum | Bacteria | 44 | 1 |
| 203692 | Spirochaetia | Class | Bacteria | 44 | 1 |
| 204432 | Terriglobia | Class | Bacteria | 31 | 1 |
| 204433 | Terriglobales | Order | Bacteria | 30 | 1 |
| 204441 | Rhodospirillales | Order | Bacteria | 143 | 4 |
| 204455 | Rhodobacterales | Order | Bacteria | 396 | 13 |
| 204457 | Sphingomonadales | Order | Bacteria | 274 | 9 |
| 204458 | Caulobacterales | Order | Bacteria | 36 | 1 |
| 206351 | Neisseriales | Order | Bacteria | 62 | 2 |
| 206389 | Rhodocyclales | Order | Bacteria | 37 | 1 |
| 213115 | Desulfovibrionales | Order | Bacteria | 39 | 1 |
| 213849 | Campylobacterales | Order | Bacteria | 78 | 2 |
| 216572 | Oscillospiraceae | Family | Bacteria | 84 | 2 |
| 335929 | Erythrobacteraceae | Family | Bacteria | 70 | 2 |
| 423349 | *Mucilaginibacter* | Genus | Bacteria | 36 | 1 |
| 563835 | Chitinophagaceae | Family | Bacteria | 68 | 2 |
| 681950 | Glomerellaceae | Family | Fungi | 22 | 1 |
| 768503 | Cytophagia | Class | Bacteria | 141 | 4 |
| 768507 | Cytophagales | Order | Bacteria | 141 | 4 |
| 909932 | Negativicutes | Class | Bacteria | 48 | 1 |
| 1028384 | Glomerellales | Order | Fungi | 22 | 1 |
| 1131492 | Aspergillaceae | Family | Fungi | 103 | 5 |
| 1643682 | Geodermatophilales | Order | Bacteria | 32 | 1 |
| 1706369 | Cellvibrionales | Order | Bacteria | 42 | 1 |
| 1737404 | Tissierellia | Class | Bacteria | 35 | 1 |
| 1737405 | Tissierellales | Order | Bacteria | 35 | 1 |
| 1775411 | Rhodanobacteraceae | Family | Bacteria | 38 | 1 |
| 1822464 | *Paraburkholderia* | Genus | Bacteria | 30 | 1 |
| 1853228 | Chitinophagia | Class | Bacteria | 68 | 2 |
| 1853229 | Chitinophagales | Order | Bacteria | 68 | 2 |
| 1853232 | Hymenobacteraceae | Family | Bacteria | 39 | 1 |
| 1866885 | *Mycolicibacterium* | Genus | Bacteria | 32 | 1 |
| 1890424 | Synechococcales | Order | Bacteria | 31 | 1 |
| 1913637 | Mucoromycota | Phylum | Fungi | 41 | 2 |
| 2691354 | Pirellulales | Order | Bacteria | 40 | 1 |
| 2726947 | Mycosphaerellales | Order | Fungi | 24 | 1 |
| 2762318 | Weeksellaceae | Family | Bacteria | 37 | 1 |
| 2854170 | Roseobacteraceae | Family | Bacteria | 196 | 6 |
| 2887326 | Moraxellales | Order | Bacteria | 34 | 1 |
| 2975441 | Sphaerotilaceae | Family | Bacteria | 39 | 1 |
| 3028117 | Cyanophyceae | Class | Bacteria | 178 | 5 |
| 3031449 | Desulfovibrionia | Class | Bacteria | 39 | 1 |
| 3031852 | Epsilonproteobacteria | Class | Bacteria | 80 | 2 |
| 3082720 | Peptostreptococcales | Order | Bacteria | 50 | 1 |
| 3085636 | Lachnospirales | Order | Bacteria | 138 | 4 |
| | | | | Total | |
| | | | | 30497 | 974 |

**Supplementary Table 3.** Detailed list of the 261 sets on which benchmark was performed. Link to the corresponding UniProt website is given for each proteome ID index.

| Set No. | Taxon name | Proteome IDs |
|---|---|---|
| 136.0 | Spirochaetales | 503, 811, 1807, 2967, 3571, 6546, 6852, 7254, 7383, 8212, 9222, 9223, 14541, 14605, 15620, 18680, 42527, 182360, 186400, 190395, 190423, 253869, 294625, 323824, 324209, 515827, 587760, 595917, 660053, 671908 |
| 237.1 | *Flavobacterium* | 5638, 6394, 30129, 30152, 31466, 93807, 184232, 184488, 198345, 198648, 216605, 237056, 237310, 244527, 244929, 244937, 245429, 245449, 247903, 271937, 289734, 296862, 297407, 316154, 557042, 609172, 625735, 630340, 675047, 1139260 |
| 265.0 | *Paracoccus* | 361, 15480, 32743, 183635, 183790, 191257, 198307, 199054, 199125, 199180, 199344, 199502, 218023, 233742, 234530, 234882, 252023, 253345, 283587, 285530, 293520, 309747, 316225, 321562, 434754, 442533, 478740, 556552, 608594, 640485 |
| 286.0 | *Pseudomonas* | 556, 686, 15503, 22611, 25241, 29493, 29499, 30063, 31652, 32531, 33462, 56648, 64137, 94259, 198407, 199524, 232228, 242957, 243063, 243359, 243778, 244064, 244119, 251762, 268166, 270661, 326483, 501989, 615613, 635983 |
| 356.12 | Hyphomicrobiales | 8952, 17669, 18524, 18542, 33187, 38011, 51058, 53043, 53675, 70354, 183447, 186364, 199236, 199466, 199667, 238563, 244780, 254764, 295097, 295238, 321085, 324738, 325303, 433101, 515290, 528286, 583454, 585507, 596427, 602124 |
| 379.0 | *Rhizobium* | 1936, 9319, 17669, 29602, 31368, 34908, 51333, 52057, 52066, 52122, 52161, 93137, 186894, 192550, 199253, 219167, 221890, 242816, 243756, 245252, 248925, 295383, 295799, 316801, 325303, 524535, 547048, 582090, 588925, 633219 |
| 433.0 | Acetobacteraceae | 245, 1176, 1963, 6375, 19250, 19760, 32668, 32679, 179145, 188604, 188879, 194822, 239724, 247565, 249065, 295023, 317078, 317214, 321405, 321746, 325255, 460715, 463253, 553193, 562254, 580654, 597459, 597507, 600101, 664073 |
| 468.0 | Moraxellaceae | 430, 546, 5740, 7477, 13117, 14568, 18415, 18418, 23785, 23788, 23795, 35860, 64939, 76238, 92616, 93391, 185895, 186001, 187495, 188357, 190435, 191094, 196536, 242317, 243900, 244223, 245977, 253940, 256774, 292423 |
| 481.0 | Neisseriaceae | 425, 535, 2768, 3009, 3019, 3344, 4088, 4105, 4207, 10457, 17813, 18554, 31390, 31392, 36027, 76625, 193118, 198238, 219669, 238938, 254209, 254293, 254651, 268229, 269923, 272771, 279284, 308891, 325536, 653156 |
| 506.0 | Alcaligenaceae | 1977, 16497, 19095, 19805, 29629, 36382, 76848, 78558, 93512, 184226, 194139, 196306, 214603, 216496, 216947, 235994, 236176, 238308, 244571, 283474, 288599, 294692, 439382, 494269, 502685, 559809, 580517, 608345, 675920, 739565 |
| 543.0 | Enterobacteriaceae | 260, 558, 625, 747, 1006, 1122, 1123, 1410, 2032, 2084, 2716, 5726, 6872, 7841, 8216, 10297, 14012, 27726, 28653, 29462, 29481, 36196, 37393, 53401, 187148, 280899, 302163, 516181, 654998, 659047 |
| 641.0 | Vibrionaceae | 537, 593, 2943, 16562, 29994, 31672, 32303, 33449, 33633, 35909, 36426, 37515, 51221, 78503, 92876, 94070, 94165, 94936, 184608, 193432, 198854, 215148, 219336, 232179, 235563, 235828, 326936, 464262, 694232, 1155586 |
| 662.0 | *Vibrio* | 584, 2493, 2943, 16562, 16567, 16895, 29994, 31672, 33673, 37515, 51221, 92876, 93173, 94070, 184608, 188276, 189475, 193432, 198854, 219336, 232179, 235563, 235828, 281112, 462621, 464262, 533102, 694232, 1139488, 1155586 |
| 815.0 | Bacteroidaceae | 1414, 2861, 3416, 6731, 7995, 14212, 17831, 17907, 17962, 18018, 18063, 18121, 18362, 18372, 18392, 18406, 18439, 56419, 92631, 95517, 95657, 184436, 195527, 196631, 262405, 264777, 279562, 284249, 324383, 651085 |
| 838.0 | *Prevotella* | 3327, 3829, 5697, 10862, 15929, 18046, 18054, 18062, 18111, 18116, 18147, 18183, 18184, 18343, 18353, 18357, 18375, 18727, 29597, 29732, 70533, 198560, 198779, 199373, 199625, 215645, 216632, 278983, 321612, 825483 |
| 976.24 | Bacteroidota | 588, 852, 3327, 7590, 8898, 17962, 18199, 32747, 75267, 76582, 187464, 198813, 199312, 199514, 236454, 241507, 244677, 249239, 249542, 253919, 255317, 263900, 266118, 270538, 286701, 295260, 297861, 317847, 437116, 503278 |
| 1117.3 | Cyanobacteriota | 268, 1423, 1566, 1961, 4344, 10475, 10478, 11180, 16960, 27395, 76925, 184315, 194922, 217698, 218238, 226442, 238762, 239001, 239589, 276103, 318453, 594557, 607397, 620559, 624057, 639877, 640021, 641405, 707356, 729701 |
| 1161.0 | Nostocales | 1191, 1511, 2483, 10378, 10380, 10390, 10474, 10475, 10477, 53372, 65745, 76925, 198313, 218238, 218621, 218785, 239589, 245124, 252906, 268857, 271624, 276103, 282767, 593846, 596768, 599391, 629098, 641405, 683511, 729701 |
| 1224.73 | Pseudomonadota | 1695, 1844, 2745, 5740, 11834, 14568, 19113, 31843, 35397, 58074, 180009, 183107, 195667, 198645, 199054, 218896, 241074, 244817, 245708, 283474, 284395, 285530, 310016, 321172, 465810, 543279, 546031, 561045, 606196, 631034 |
| 1236.27 | Gammaproteobacteria | 1122, 1947, 6286, 10320, 32414, 35860, 54457, 55136, 59419, 61457, 70250, 92671, 181743, 198575, 198706, 218896, 234640, 235547, 242642, 253740, 254575, 281112, 282106, 302163, 310227, 316365, 321272, 501602, 596063, 635983 |
| 1239.36 | Bacillota | 3195, 3330, 4069, 6316, 7523, 11124, 11912, 17747, 18136, 18364, 30408, 30832, 32409, 37233, 51230, 51657, 51813, 78534, 95380, 195141, 199476, 217785, 288024, 323257, 502248, 562464, 595468, 656813, 679779, 681343 |

**Supplementary Table 3.** (Continued.)

| Set No. | Taxon name | Proteome IDs |
|---|---|---|
| 1268.2 | Micrococcaceae | 30301, 30982, 53171, 60433, 95776, 177882, 190514, 192359, 219947, 247286, 277032, 280861, 283134, 292685, 295411, 305233, 307000, 325957, 326852, 460157, 516404, 516421, 544090, 600171, 633136, 638848, 655366, 664164, 676885, 1139502 |
| 1300.0 | Streptococcaceae | 585, 586, 750, 2196, 2814, 2815, 3330, 7946, 8520, 9077, 70387, 70638, 72050, 182508, 186437, 198604, 218181, 218689, 242246, 254082, 254634, 254924, 269148, 269374, 318879, 439550, 522720, 562464, 563349, 644875 |
| 1301.0 | *Streptococcus* | 449, 585, 586, 750, 1170, 1452, 2148, 2814, 2815, 4514, 5388, 7946, 70387, 70638, 72050, 182508, 186437, 198604, 249495, 254082, 254634, 254924, 269148, 279194, 281771, 291494, 522720, 563349, 644875, 660801 |
| 1385.16 | Bacillales | 3445, 6315, 8816, 17973, 18296, 27985, 28863, 35996, 78534, 92623, 180019, 180199, 189761, 198734, 198936, 215059, 248066, 258625, 265711, 270468, 275473, 282076, 295689, 437773, 515679, 593626, 618460, 676456, 829401, 1084197 |
| 1386.1 | *Bacillus* | 606, 27602, 27822, 36953, 60713, 67625, 95209, 180019, 180199, 185615, 199133, 199578, 216230, 219546, 226441, 253314, 258625, 263943, 265711, 269701, 325032, 437773, 472971, 481043, 531594, 575961, 666543, 678477, 678491, 679638 |
| 1485.2 | *Clostridium* | 818, 1986, 2730, 8937, 10420, 17118, 17809, 17939, 18022, 18023, 18136, 18268, 19426, 30009, 75531, 76603, 175744, 184869, 192468, 220840, 254664, 260680, 286268, 287890, 306888, 320543, 545014, 559354, 580568, 594441 |
| 1653.1 | Corynebacteriaceae | 545, 582, 1473, 2198, 4208, 6078, 10445, 14809, 15388, 16943, 17152, 23703, 29914, 35199, 70591, 76947, 177657, 182237, 185479, 217209, 238368, 244989, 278422, 320791, 424462, 516320, 594681, 645966, 658613, 762474 |
| 1663.0 | *Arthrobacter* | 30301, 53253, 54460, 176251, 181917, 190514, 195913, 199258, 237061, 239972, 244710, 247980, 273807, 280861, 283134, 295411, 295511, 296209, 305124, 306826, 317144, 326852, 543556, 544090, 547458, 664164, 676885, 683653, 687525, 692783 |
| 1678.0 | *Bifidobacterium* | 439, 3191, 8702, 28730, 29004, 29014, 29033, 29046, 29050, 29052, 29060, 29067, 29096, 29108, 30636, 216352, 216444, 216451, 229095, 229239, 242610, 252530, 287470, 287533, 287609, 288052, 326336, 441772, 482084, 718821 |
| 1716.0 | *Corynebacterium* | 492, 4218, 10445, 14809, 15388, 17152, 19222, 33457, 35368, 50488, 60016, 70591, 76947, 177600, 177657, 199350, 221653, 244989, 247696, 251577, 254467, 278422, 312032, 317518, 326711, 439070, 515743, 516320, 645966, 762474 |
| 1760.24 | Actinomycetes | 5063, 7076, 10411, 16033, 23703, 31501, 50821, 51005, 51073, 94243, 175826, 184452, 185596, 237697, 254869, 262621, 281230, 292235, 319746, 321701, 323454, 425099, 431080, 473525, 537260, 541033, 581206, 606172, 653493, 694501 |
| 1762.0 | Mycobacteriaceae | 757, 1020, 1419, 1584, 9224, 36176, 36334, 51677, 70146, 70612, 77143, 94224, 192566, 192801, 193090, 193564, 220914, 268285, 279306, 282551, 295165, 320513, 323700, 430146, 466445, 466517, 467006, 467105, 467130, 1064782 |
| 1763.0 | *Mycobacterium* | 1020, 1190, 1419, 1584, 1988, 25947, 36334, 51677, 53444, 70146, 77143, 92025, 92027, 92060, 92151, 93629, 94224, 94976, 179734, 192566, 193990, 194000, 223528, 320513, 465361, 466396, 467105, 467130, 554965, 682202 |
| 1817.0 | *Nocardia* | 6304, 6820, 17048, 19150, 19401, 76512, 188836, 215506, 219565, 246410, 252586, 254869, 255355, 255467, 266677, 279275, 294615, 429627, 431401, 432464, 438448, 503540, 515512, 540412, 540698, 565715, 586827, 612956, 638263, 655751 |
| 1839.0 | *Nocardioides* | 15971, 33772, 50886, 51395, 51495, 51699, 77868, 198649, 198832, 199004, 246018, 279994, 283644, 291189, 294853, 304990, 307087, 308550, 324351, 325003, 326325, 433406, 473325, 502035, 530424, 535511, 537326, 549911, 616839, 640489 |
| 1873.0 | *Micromonospora* | 1908, 3448, 10307, 32254, 70620, 186004, 186700, 198210, 198217, 198251, 198765, 198937, 199001, 199393, 199408, 199413, 199699, 240342, 242415, 258607, 262621, 277671, 281230, 281726, 283832, 285744, 295555, 427831, 478148, 578819 |
| 1883.4 | *Streptomyces* | 2357, 4217, 9036, 10411, 14390, 14410, 52945, 53429, 78145, 94479, 186200, 189065, 189443, 199155, 217446, 218662, 235945, 265325, 281963, 294981, 326553, 472335, 481583, 501663, 637788, 645555, 656732, 677732, 689511, 693098 |
| 2004.0 | Streptosporangiaceae | 6640, 18869, 77701, 190797, 198923, 198953, 199361, 236732, 249304, 253094, 253303, 284824, 286861, 295157, 306628, 308705, 530928, 540685, 555564, 562352, 568380, 578449, 586042, 605992, 610966, 640052, 645217, 655287, 674234, 1143474 |
| 2012.0 | Thermomonosporaceae | 183413, 198318, 236723, 242367, 251891, 256661, 261811, 262882, 272400, 282674, 288610, 294513, 295578, 295596, 295623, 316096, 316706, 323380, 432015, 468735, 487268, 501240, 539313, 546324, 572680, 579250, 614047, 669179, 1165074, 1165124 |
| 2037.0 | Actinomycetales | 3045, 3994, 5351, 6742, 9888, 10301, 13015, 18843, 54404, 78368, 176288, 182744, 185628, 186785, 245283, 250192, 266895, 270021, 271272, 273303, 276899, 280344, 284680, 293036, 594637, 595895, 595973, 614239, 627538, 683503 |
| 2049.0 | Actinomycetaceae | 2941, 3045, 3822, 5351, 6742, 9888, 10301, 16495, 54404, 78368, 176288, 176960, 184291, 185612, 185628, 186785, 250192, 266895, 269542, 271272, 273303, 276899, 280444, 284680, 293036, 594637, 595973, 614239, 627538, 683503 |

**Supplementary Table 3.** (Continued.)

| Set No. | Taxon name | Proteome IDs |
|---|---|---|
| 2062.2 | Streptomycetaceae | 4184, 8043, 14629, 28341, 37702, 37982, 51136, 54011, 217789, 237426, 240682, 245455, 250268, 271004, 276963, 295370, 317940, 435531, 450869, 472335, 473262, 481552, 509480, 509509, 528419, 600365, 603708, 608024, 630936, 655443 |
| 2070.0 | Pseudonocardiaceae | 328, 5087, 6281, 7809, 13968, 19277, 31419, 33409, 62973, 184363, 185596, 186040, 192591, 194360, 198348, 198582, 198967, 199494, 199614, 204221, 238362, 239494, 249915, 274515, 295444, 325787, 478898, 550714, 650035, 660680 |
| 4890.0 | Ascomycota | 36893, 55045, 184073, 186583, 190274, 231358, 240493, 241462, 248349, 248423, 266453, 544331, 582016, 738349, 774326, 788993, 800041, 813383, 1140510, 1152130 |
| 4890.1 | Ascomycota | 6701, 16922, 30104, 182235, 191612, 244722, 256690, 298493, 315783, 326289, 326340, 567885, 569620, 631181, 664169, 664534, 756132, 799441, 811619, 1140502 |
| 4890.10 | Ascomycota | 8062, 8181, 8513, 16923, 27730, 30663, 37505, 184546, 222788, 244309, 246702, 249402, 266272, 452235, 541154, 558688, 694255, 800092, 813426, 813461 |
| 4890.11 | Ascomycota | 1197, 16801, 19471, 28524, 30706, 53411, 54466, 95085, 295703, 297299, 305883, 310108, 319663, 326198, 554235, 654913, 676310, 736672, 799772, 829364 |
| 4890.12 | Ascomycota | 1067, 9058, 54771, 223968, 237438, 241587, 241690, 243732, 284375, 288168, 325902, 326931, 327118, 434172, 515153, 748025, 782241, 800039, 1147733, 1147747 |
| 4890.13 | Ascomycota | 2762, 7115, 7703, 16932, 19375, 20467, 31176, 53328, 76632, 77069, 91967, 243015, 249363, 249829, 325579, 758155, 800038, 809789, 1147760, 1149163 |
| 4890.14 | Ascomycota | 709, 8066, 11761, 15100, 16935, 34112, 50424, 70168, 190831, 191342, 297527, 326757, 398389, 578531, 737391, 758603, 799539, 800040, 1056384, 1146351 |
| 4890.15 | Ascomycota | 314, 598, 1471, 5627, 9096, 19484, 54821, 76881, 94455, 218381, 234585, 285146, 593570, 664203, 664521, 697127, 777438, 799424, 799537, 1152049 |
| 4890.16 | Ascomycota | 2037, 53617, 76837, 78340, 78544, 95728, 191408, 191691, 193144, 193689, 194361, 237481, 249293, 325492, 481858, 694050, 701801, 813444, 1149165, 1150942 |
| 4890.17 | Ascomycota | 689, 1610, 1631, 1881, 1997, 5220, 6702, 37136, 37696, 54302, 76874, 189513, 192386, 219602, 235786, 325672, 509704, 700596, 749293, 785799 |
| 4890.18 | Ascomycota | 1640, 2499, 8784, 9097, 54544, 70121, 76552, 184300, 191518, 224080, 226031, 265631, 266234, 285775, 286921, 326924, 613401, 696573, 722485, 1056012 |
| 4890.19 | Ascomycota | 560, 1745, 2311, 5222, 7796, 9882, 24533, 94112, 177622, 237631, 243797, 248340, 250266, 596902, 637239, 696280, 720189, 799777, 1150879, 1150941 |
| 4890.2 | Ascomycota | 267, 1261, 2258, 2866, 8536, 19487, 37904, 70720, 78343, 178129, 190312, 241107, 242519, 253071, 275385, 319160, 510647, 606974, 799291, 799423 |
| 4890.20 | Ascomycota | 6310, 6911, 7304, 8142, 15530, 16934, 19478, 54516, 70133, 214365, 225277, 233524, 235728, 245910, 246991, 247810, 248961, 504637, 531561, 800036 |
| 4890.21 | Ascomycota | 591, 2038, 2530, 27920, 42958, 77535, 94389, 184383, 241818, 247233, 253472, 509510, 562929, 572817, 604273, 730381, 799429, 799437, 827724, 1147746 |
| 4890.22 | Ascomycota | 599, 2036, 7322, 19804, 54304, 70501, 76584, 183809, 192596, 230249, 236621, 316270, 672032, 730481, 790833, 799324, 799436, 799438, 799771, 1153618 |
| 4890.23 | Ascomycota | 2428, 5426, 16936, 18001, 30651, 94801, 184304, 215289, 235371, 238813, 266152, 275078, 277212, 292447, 308549, 325780, 624404, 769157, 769528, 824596 |
| 4890.24 | Ascomycota | 1805, 7978, 18087, 53342, 92555, 95038, 193240, 235672, 238350, 252748, 253729, 286045, 286134, 290900, 622797, 635477, 654918, 756346, 1148299, 1152087 |
| 4890.25 | Ascomycota | 1798, 1996, 6753, 16928, 30106, 30143, 53029, 70700, 234275, 287544, 297452, 309340, 326532, 605986, 626568, 641853, 781932, 830671, 1140562, 1147752 |
| 4890.3 | Ascomycota | 5206, 5666, 6790, 8984, 16800, 19376, 30641, 34291, 54342, 242791, 326565, 326799, 431533, 547976, 750502, 756921, 813427, 1149074, 1149079, 1152649 |
| 4890.4 | Ascomycota | 724, 5221, 6039, 8782, 11096, 14480, 53259, 76744, 78397, 94526, 94569, 184188, 240883, 243723, 574317, 699042, 799770, 837801, 1152300, 1152885 |
| 4890.5 | Ascomycota | 2035, 2668, 8673, 30672, 54383, 182334, 184356, 191024, 191285, 226192, 241546, 242955, 276215, 283383, 297280, 310066, 660729, 701341, 717696, 1154252 |
| 4890.6 | Ascomycota | 15441, 16931, 27002, 77248, 184330, 188318, 236664, 243498, 254866, 283841, 283895, 326950, 532311, 653565, 661057, 714618, 799757, 799764, 1141434, 1147782 |
| 4890.7 | Ascomycota | 559, 707, 2497, 16933, 30753, 30816, 36947, 53841, 75602, 78576, 184499, 194280, 256645, 276864, 277580, 297229, 326268, 504636, 630445, 800035 |
| 4890.8 | Ascomycota | 2669, 19473, 33647, 94444, 179179, 182658, 186955, 191672, 267821, 283090, 287144, 297716, 307173, 546213, 799750, 799767, 800082, 800094, 800235, 1149954 |
| 4890.9 | Ascomycota | 1294, 6706, 9328, 27238, 53599, 94285, 176998, 196158, 215305, 234474, 517252, 536711, 557566, 593566, 639643, 651452, 799439, 800096, 801428, 1150904 |
| 4891.0 | Saccharomycetes | 559, 1997, 2311, 2866, 7703, 94112, 94285, 94801, 95085, 95728, 189513, 190274, 190831, 244309, 290900, 398389, 510647, 662931, 769528, 837801 |
| 4891.1 | Saccharomycetes | 591, 599, 707, 709, 1640, 1996, 5221, 5222, 19375, 92555, 94389, 95038, 238350, 249293, 253472, 307173, 697127, 769157, 790833, 1152885 |
| 4891.2 | Saccharomycetes | 598, 1300, 2036, 2037, 2428, 5220, 5627, 5666, 6310, 9328, 94455, 182334, 191024, 196158, 230249, 285775, 286412, 292447, 509704, 774326 |

**Supplementary Table 3.** (Continued.)

| Set No. | Taxon name | Proteome IDs |
|---|---|---|
| 4892.0 | Saccharomycetales | 689, 707, 2037, 5220, 6310, 8673, 9328, 19375, 94112, 94285, 94801, 189513, 196158, 249293, 253472, 398389, 509704, 662931, 694255, 774326 |
| 4892.1 | Saccharomycetales | 314, 591, 599, 1300, 1997, 2311, 5221, 5222, 5627, 8536, 92555, 95085, 182334, 190274, 285775, 286412, 292447, 769157, 769528, 837801 |
| 4892.2 | Saccharomycetales | 559, 709, 1640, 2036, 2258, 2428, 5666, 6790, 7703, 54304, 94389, 94455, 95038, 191024, 230249, 244309, 307173, 510647, 697127, 788993 |
| 4893.0 | Saccharomycetaceae | 267, 591, 689, 1640, 2036, 2311, 2428, 2866, 5220, 5627, 5666, 6310, 8536, 19375, 54304, 190274, 190831, 196158, 509704, 510647 |
| 5042.0 | Eurotiales | 9882, 18001, 30143, 37505, 179179, 184073, 184383, 184499, 186955, 190312, 191691, 249402, 325579, 630445, 637239, 701341, 1147746, 1147752, 1150879, 1150942 |
| 5042.1 | Eurotiales | 1294, 1745, 55045, 184188, 184304, 184546, 191342, 191408, 191672, 247810, 249829, 319663, 326289, 326532, 326950, 509510, 641853, 653565, 1147782, 1149074 |
| 5042.2 | Eurotiales | 6702, 36893, 70168, 94569, 191518, 215289, 234275, 234585, 248961, 249363, 256690, 283841, 286921, 325672, 326565, 631181, 1148299, 1149165, 1150941, 1153618 |
| 5042.3 | Eurotiales | 560, 6706, 19376, 37696, 42958, 214365, 215305, 231358, 246702, 248340, 248423, 253729, 325780, 326931, 541154, 1146351, 1149079, 1149954, 1150904, 1154252 |
| 5042.4 | Eurotiales | 724, 2530, 6701, 19804, 30104, 34291, 54383, 177622, 184300, 188318, 191612, 218381, 234474, 247233, 326268, 326799, 327118, 1147733, 1147760, 1149163 |
| 5052.0 | *Aspergillus* | 6701, 19804, 36893, 37505, 94569, 184499, 184546, 188318, 215289, 231358, 234275, 234474, 246702, 249402, 256690, 286921, 326532, 452235, 630445, 637239 |
| 5052.1 | *Aspergillus* | 6706, 34291, 54771, 184300, 184304, 184356, 215305, 248340, 248423, 248961, 253729, 325579, 325672, 325780, 326799, 326931, 327118, 541154, 641853, 661057 |
| 5073.0 | *Penicillium* | 9882, 30143, 42958, 55045, 186955, 191285, 191612, 191672, 218381, 631181, 701341, 1147733, 1147746, 1147747, 1147760, 1147782, 1149163, 1149954, 1150879, 1153618 |
| 5073.1 | *Penicillium* | 724, 19376, 30104, 37696, 70168, 177622, 191408, 191518, 191691, 1141434, 1146351, 1147752, 1148299, 1149074, 1149079, 1150904, 1150941, 1150942, 1152649, 1154252 |
| 5125.0 | Hypocreales | 5206, 7115, 7978, 16801, 27002, 28524, 36947, 78397, 78544, 236664, 532311, 557566, 562929, 567885, 569620, 730481, 737391, 813383, 813427, 824596 |
| 5125.1 | Hypocreales | 30106, 54544, 76744, 219602, 240493, 252748, 265631, 287544, 582016, 622797, 626568, 635477, 694050, 696573, 717696, 748025, 750502, 782241, 813444, 1152130 |
| 5125.2 | Hypocreales | 1610, 2499, 8984, 30663, 31176, 50424, 70720, 76874, 236621, 243498, 245910, 266272, 287144, 574317, 593570, 720189, 722485, 736672, 1140502, 1152087 |
| 5125.3 | Hypocreales | 9097, 16800, 34112, 37904, 76580, 76881, 78340, 226192, 241546, 241587, 253071, 277212, 536711, 544331, 554235, 558688, 604273, 738349, 777438, 829364 |
| 5125.4 | Hypocreales | 2762, 5426, 9096, 16928, 30753, 37136, 91967, 235728, 237481, 241690, 266152, 266234, 315783, 517252, 547976, 605986, 749293, 811619, 827724, 1152049 |
| 5178.0 | Helotiales | 1798, 178129, 184330, 235371, 235672, 235786, 238813, 241818, 242519, 254866, 256645, 297280, 297299, 297452, 326757, 531561, 624404, 696280, 701801, 1152300 |
| 5204.0 | Basidiomycota | 561, 2149, 53477, 54097, 62823, 76761, 76871, 78595, 184267, 198372, 245768, 249464, 256964, 284706, 294933, 305067, 307440, 521943, 629468, 1063166 |
| 5204.1 | Basidiomycota | 1072, 1861, 6352, 14071, 53257, 53890, 53989, 77086, 193218, 194127, 199069, 218811, 235388, 242287, 250043, 279236, 287166, 308652, 313359, 759537 |
| 5204.2 | Basidiomycota | 5242, 17559, 53392, 53424, 53558, 53611, 53819, 54166, 54477, 92583, 92666, 193986, 217199, 305948, 324022, 623687, 664032, 714275, 807025, 1142393 |
| 5204.3 | Basidiomycota | 17200, 30669, 54549, 76154, 76842, 77521, 94020, 218334, 219338, 310668, 324748, 567179, 609673, 703269, 772434, 807353, 807769, 813600, 815604, 886653 |
| 5204.4 | Basidiomycota | 8493, 16930, 30653, 53712, 54270, 54485, 77684, 92730, 186303, 237144, 245783, 246740, 298030, 323386, 365756, 559027, 736335, 765509, 812287, 813824 |
| 5204.5 | Basidiomycota | 7431, 9131, 15241, 27361, 54564, 76727, 76738, 94043, 94065, 183567, 193067, 217790, 249723, 284842, 322225, 322482, 719766, 724874, 807469, 1049176 |
| 5204.6 | Basidiomycota | 27073, 27195, 27222, 27265, 30671, 54007, 54018, 54538, 54988, 76532, 78113, 245771, 311382, 320762, 521872, 799118, 807306, 807342, 823399, 1150266 |
| 5338.0 | Agaricales | 27073, 27222, 53424, 54549, 76154, 242287, 283269, 284706, 284842, 298030, 307440, 567179, 629468, 724874, 807025, 807342, 807469, 812287, 1049176, 1150266 |
| 5338.1 | Agaricales | 1861, 8493, 53712, 54007, 54270, 54477, 54988, 218334, 305067, 308652, 320762, 521872, 521943, 623687, 664032, 772434, 807353, 813824, 1063166, 1142393 |
| 5455.0 | *Colletotrichum* | 8782, 11096, 14480, 15530, 20467, 27238, 70121, 76552, 76584, 186583, 295703, 305883, 326340, 434172, 613401, 639643, 654918, 699042, 781932, 830671 |
| 5506.0 | *Fusarium* | 9096, 9097, 16800, 30753, 37904, 91967, 219602, 241587, 266152, 277212, 536711, 546213, 547976, 569620, 605986, 622797, 736672, 737391, 813427, 1152087 |
| 5506.1 | *Fusarium* | 5206, 30663, 70720, 236664, 245910, 265631, 266234, 287144, 288168, 532311, 544331, 558688, 574317, 582016, 593570, 604273, 635477, 730481, 750502, 1140502 |

**Supplementary Table 3.** (Continued.)

| Set No. | Taxon name | Proteome IDs |
|---|---|---|
| 13687.1 | *Sphingomonas* | 33200, 50871, 51371, 51455, 51642, 51854, 51880, 185161, 188729, 194475, 198281, 198824, 198912, 199586, 218151, 218323, 241167, 244162, 266693, 286100, 286883, 315673, 321250, 436053, 515367, 529795, 558192, 613564, 623067, 1139410 |
| 28056.2 | Micromonosporaceae | 235, 3448, 5440, 53244, 186004, 197174, 198251, 198362, 199408, 199699, 242415, 287606, 295555, 482800, 542742, 578112, 578819, 587527, 598146, 619260, 623608, 629619, 635606, 649753, 660611, 676360, 680286, 680866, 1143480, 1165079 |
| 28211.24 | Alphaproteobacteria | 1302, 1591, 5713, 8850, 17808, 28638, 35287, 51777, 53043, 53455, 184221, 198703, 198912, 231259, 237344, 244932, 254889, 284322, 317894, 320653, 471755, 530268, 533469, 538147, 546701, 547048, 551327, 554286, 609121, 1144323 |
| 28216.16 | Betaproteobacteria | 7883, 8392, 51759, 63254, 77549, 184226, 186110, 197446, 197468, 198781, 199646, 229897, 237839, 239406, 241193, 247811, 266302, 288813, 295525, 297839, 301751, 305539, 318422, 325536, 444316, 487350, 515986, 559809, 664731, 681964 |
| 28256.0 | Halomonadaceae | 4512, 14463, 19113, 63387, 77875, 179615, 184346, 187327, 190911, 198500, 198641, 198654, 198693, 199040, 199677, 218896, 219993, 235346, 267342, 276016, 287023, 321272, 321275, 327197, 437638, 448235, 487929, 501340, 518892, 595089 |
| 29547.1 | Campylobacterota | 422, 760, 2714, 5085, 7032, 7727, 8139, 15520, 18143, 29714, 92701, 92884, 194141, 194309, 199227, 216568, 217944, 229442, 230780, 233248, 240535, 251062, 253850, 255139, 256695, 256842, 290191, 502820, 503264, 593836 |
| 31953.0 | Bifidobacteriaceae | 439, 2456, 5777, 6173, 8693, 16519, 29004, 29033, 29046, 29050, 29052, 29055, 29067, 29080, 216352, 216444, 216451, 216454, 229095, 229239, 231451, 242610, 243657, 252530, 287533, 287609, 326336, 441772, 469292, 482084 |
| 31957.0 | Propionibacteriaceae | 936, 7947, 14417, 93501, 184512, 187142, 188235, 188324, 188342, 199475, 215896, 216300, 216311, 252770, 273044, 274803, 275256, 277858, 291933, 292373, 295371, 316196, 317638, 319263, 425099, 435304, 523079, 527616, 569914, 613840 |
| 31979.3 | Clostridiaceae | 1325, 3081, 8937, 17118, 17991, 18047, 18076, 18120, 30009, 50326, 52015, 76603, 77407, 175744, 184035, 184526, 190080, 190748, 198524, 198597, 199127, 239471, 286268, 286369, 304999, 306888, 308489, 324646, 664218, 724672 |
| 31989.2 | Paracoccaceae | 4507, 30004, 50471, 183002, 186098, 194664, 198417, 199125, 201838, 207598, 215377, 215760, 230388, 234539, 237655, 244810, 244932, 245680, 279673, 282002, 293520, 316225, 442533, 478183, 484076, 572377, 598196, 619033, 619079, 638981 |
| 32003.1 | Nitrosomonadales | 383, 1235, 1629, 2440, 2718, 2743, 5019, 12179, 31637, 56322, 70578, 77545, 77549, 185062, 195558, 198620, 198729, 198764, 198772, 198814, 199561, 199646, 245081, 248024, 275137, 295135, 316473, 316827, 501534, 681964 |
| 32033.1 | Xanthomonadaceae | 1890, 5870, 8632, 29385, 30017, 30518, 33067, 50805, 51386, 51738, 51802, 61569, 218968, 219374, 239898, 249447, 251842, 262917, 269708, 289784, 291286, 291562, 315167, 316584, 317199, 319980, 322165, 519004, 639274, 680299 |
| 33882.0 | *Microbacterium* | 24001, 28883, 33900, 34098, 51381, 93355, 187047, 196320, 199183, 199422, 215463, 245623, 252505, 253508, 273158, 297593, 298650, 318564, 320235, 321196, 321225, 325827, 326838, 422989, 432812, 440939, 526083, 537775, 633205, 657592 |
| 33958.2 | Lactobacillaceae | 432, 1652, 1991, 4069, 4115, 4567, 5444, 17248, 18559, 37778, 50911, 50969, 51412, 51461, 51638, 51639, 51672, 51697, 51733, 51845, 51931, 51992, 52012, 52013, 94714, 199376, 290602, 294321, 295257, 1139006 |
| 41297.5 | Sphingomonadaceae | 4728, 11816, 13072, 37930, 77614, 188729, 194475, 218366, 238954, 253918, 266860, 276254, 282837, 286100, 286883, 287772, 298213, 319092, 326488, 426325, 509198, 546701, 552700, 552757, 554342, 570166, 571950, 575068, 578569, 676996 |
| 44249.2 | *Paenibacillus* | 3445, 3900, 7523, 28123, 29409, 29453, 29512, 32320, 36611, 37688, 76927, 92024, 92663, 93309, 187480, 190188, 199177, 215509, 244407, 252415, 279446, 293568, 295418, 307943, 325218, 450917, 490800, 502136, 608782, 676098 |
| 49546.0 | Flavobacteriaceae | 3023, 3053, 3730, 5938, 8297, 32229, 45051, 93510, 93807, 94750, 182510, 184516, 195470, 198521, 198999, 219193, 219559, 244937, 245429, 248054, 256629, 276603, 290889, 295479, 321734, 321790, 535020, 761667, 1139260, 1143545 |
| 57723.0 | Acidobacteriota | 343, 2207, 2432, 6056, 6844, 7113, 31518, 182427, 198356, 199024, 236728, 253606, 269669, 279598, 289437, 290253, 292958, 295210, 321820, 513480, 515312, 515460, 515496, 540989, 568106, 589520, 593892, 647241, 648801, 1059380 |
| 69277.1 | Phyllobacteriaceae | 3250, 4291, 6786, 50665, 50670, 54487, 70107, 94412, 179771, 182840, 191905, 199768, 215931, 218091, 235507, 238563, 242763, 244404, 278398, 281647, 293719, 295131, 312852, 317542, 323300, 595886, 630142, 636264, 643405, 1149009 |
| 72274.1 | Pseudomonadales | 233, 556, 686, 2515, 15503, 16560, 30980, 32239, 33462, 63953, 64137, 68210, 94259, 184497, 185766, 198407, 198706, 199290, 231409, 232228, 243359, 244119, 270661, 278332, 280615, 315235, 327522, 463138, 476718, 615613 |
| 72275.0 | Alteromonadaceae | 6263, 6327, 9282, 11864, 19276, 35062, 37600, 53586, 56090, 68447, 70299, 175691, 176037, 184520, 219285, 233828, 244441, 245728, 247426, 256561, 275281, 291390, 293649, 298049, 317204, 601768, 606935, 631300, 664654, 664717 |

**Supplementary Table 3.** (Continued.)

| Set No. | Taxon name | Proteome IDs |
|---|---|---|
| 72293.0 | Helicobacteraceae | 422, 429, 1522, 2495, 5085, 7032, 7934, 8387, 18143, 18688, 18731, 29707, 29714, 29920, 92701, 92884, 216091, 216568, 218502, 249746, 251062, 251881, 255139, 256379, 256421, 256695, 256765, 256842, 257067, 319322 |
| 74201.0 | Verrucomicrobiota | 1031, 4837, 7013, 71392, 78486, 95228, 199432, 217265, 239907, 244023, 306196, 315648, 321577, 475117, 478417, 500777, 501812, 525652, 526501, 546464, 557872, 590740, 600139, 603141, 617628, 642829, 644507, 658278, 676169, 825051 |
| 75682.0 | Oxalobacteraceae | 329, 5089, 27604, 28843, 31572, 34315, 51728, 51760, 56897, 190838, 197535, 198284, 199391, 214937, 228593, 240505, 241421, 244112, 265955, 274350, 285190, 294829, 297258, 298438, 444318, 451565, 484015, 502415, 534388, 622890 |
| 76892.0 | Caulobacteraceae | 1816, 1868, 2696, 17808, 17812, 17826, 17837, 50923, 51150, 51182, 51604, 56905, 77603, 195546, 228945, 247763, 249254, 249524, 249725, 249842, 317223, 320100, 431269, 482044, 545037, 548978, 566663, 622580, 662572, 663918 |
| 80840.1 | Burkholderiales | 5045, 16497, 27020, 27834, 29629, 50580, 51403, 51759, 51826, 72741, 214603, 216607, 243193, 252174, 274884, 291078, 298438, 439382, 444316, 502415, 521868, 529637, 542125, 574067, 580517, 588051, 623397, 627205, 675920, 739565 |
| 80864.0 | Comamonadaceae | 374, 2482, 3856, 4811, 7938, 8332, 51275, 51403, 51912, 53300, 70169, 72741, 94760, 185680, 190750, 194358, 194432, 198552, 199369, 199766, 216607, 239709, 297564, 316798, 464549, 501978, 530032, 541185, 623043, 664731 |
| 81852.0 | Enterococcaceae | 1415, 8456, 10296, 12675, 13782, 14127, 15961, 17415, 18126, 95256, 181884, 182077, 182149, 188246, 195141, 195918, 196151, 236214, 286773, 287101, 287857, 288028, 288197, 290567, 298615, 321556, 500890, 622610, 637757, 674938 |
| 82115.0 | Rhizobiaceae | 813, 1054, 1596, 1936, 1976, 9319, 10792, 17669, 28181, 29602, 31368, 34908, 51333, 52122, 52161, 52167, 186894, 241247, 248925, 264280, 295238, 295383, 316801, 439983, 547048, 582090, 588925, 663351, 663401, 680848 |
| 84566.0 | Sphingobacteriaceae | 310, 3664, 36014, 50543, 186720, 198670, 198785, 199072, 199226, 199666, 238034, 248198, 253209, 268007, 284120, 286701, 291117, 293331, 293347, 307244, 308196, 310477, 323829, 462014, 505355, 515877, 538158, 601292, 619078, 662074 |
| 84567.0 | *Pedobacter* | 852, 3664, 28007, 36014, 50543, 71561, 78459, 94313, 192756, 198039, 198850, 199572, 199666, 240912, 244301, 245379, 248198, 284120, 291117, 293347, 295499, 295620, 295668, 307244, 308181, 309488, 310477, 601055, 601292, 651668 |
| 84998.0 | Coriobacteriia | 333, 954, 960, 2026, 3446, 3560, 4830, 6069, 6851, 12651, 14204, 16638, 18266, 31121, 54078, 70675, 182975, 190585, 195399, 195781, 195889, 253805, 269591, 273154, 321063, 400317, 468668, 486032, 503297, 681649 |
| 85004.0 | Bifidobacteriales | 439, 2456, 3191, 5777, 6173, 8693, 8702, 28730, 29014, 29033, 29080, 29082, 29093, 29108, 30636, 216352, 216444, 216451, 228755, 229095, 231451, 240228, 242610, 252530, 287533, 288052, 326336, 482084, 619536, 718821 |
| 85006.8 | Micrococcales | 12015, 16605, 33448, 35721, 50966, 51694, 53012, 57181, 195787, 218598, 241203, 253508, 265742, 273807, 276333, 278440, 297447, 297593, 297903, 298433, 315628, 316747, 321379, 516154, 540191, 540568, 592181, 616114, 619453, 1142462 |
| 85007.2 | Mycobacteriales | 1419, 4245, 4915, 9235, 10988, 11723, 16943, 19150, 35199, 44872, 51127, 51962, 92027, 183810, 186218, 194000, 199350, 279306, 294615, 427071, 431401, 467105, 467130, 467249, 475545, 503540, 540698, 565715, 641514, 1140293 |
| 85008.2 | Micromonosporales | 235, 32254, 182486, 186004, 198210, 198217, 198362, 198415, 199001, 199645, 248749, 283832, 285744, 295555, 478148, 482800, 542742, 546162, 590749, 598146, 599074, 608890, 612808, 612899, 622552, 649753, 652013, 653674, 680286, 681340 |
| 85009.2 | Propionibacteriales | 936, 50971, 51640, 184512, 187142, 188235, 194147, 198649, 198983, 199034, 199475, 216311, 275256, 279994, 288271, 291101, 293071, 293291, 294853, 295388, 308193, 313231, 325003, 326325, 523079, 527616, 530424, 616839, 662940, 683575 |
| 85010.0 | Pseudonocardiales | 7809, 33393, 62082, 184501, 185596, 186040, 198348, 199614, 204221, 238362, 249915, 272729, 282084, 292003, 294257, 294911, 295560, 295680, 321328, 323946, 501808, 533598, 535890, 552097, 569329, 580474, 597989, 660680, 1143463, 1165136 |
| 85011.4 | Kitasatosporales | 10931, 14390, 34196, 53024, 53127, 54804, 175829, 176087, 181951, 182206, 186318, 189065, 192930, 198750, 222531, 230407, 245372, 247634, 277533, 281963, 292452, 325849, 444212, 481022, 608955, 617734, 641386, 653493, 675554, 677896 |
| 85012.0 | Streptosporangiales | 31675, 198683, 198923, 198953, 215005, 238312, 249304, 253303, 294513, 295596, 306628, 309033, 316096, 501240, 523007, 534286, 576393, 579153, 579250, 583800, 586042, 605992, 614047, 628304, 640052, 644610, 646523, 660745, 669179, 1165074 |
| 85015.0 | Nocardioidaceae | 640, 3111, 50886, 50971, 51395, 51414, 51542, 51699, 51702, 77868, 194147, 198832, 199034, 199092, 230842, 244867, 246018, 275225, 276542, 293071, 293291, 307087, 308193, 309647, 500693, 530424, 549911, 640489, 662940, 663365 |
| 85016.0 | Cellulomonadaceae | 485, 849, 8460, 19753, 29839, 54314, 54319, 185663, 199404, 231693, 245142, 269289, 273256, 276333, 293764, 296469, 315994, 321181, 321386, 321484, 321720, 321723, 321798, 374843, 562124, 581206, 612875, 632740, 642125, 664209 |

**Supplementary Table 3.** (Continued.)

| Set No. | Taxon name | Proteome IDs |
|---|---|---|
| 85021.0 | Intrasporangiaceae | 5063, 8914, 13167, 19489, 19494, 30011, 30013, 35720, 35721, 51641, 76976, 175826, 182938, 199019, 233781, 237822, 278440, 316747, 317893, 319514, 319516, 321534, 321793, 431092, 437955, 554054, 573599, 588586, 592181, 677016 |
| 85023.3 | Microbacteriaceae | 24001, 33572, 33836, 37934, 50802, 50821, 51216, 51768, 51858, 69933, 190857, 195931, 199183, 228758, 238650, 243284, 243784, 270299, 274391, 275048, 298424, 321196, 422989, 501494, 515934, 517724, 573729, 610303, 636956, 1144396 |
| 85025.0 | Nocardiaceae | 4245, 6820, 8190, 13525, 19150, 19401, 76180, 76512, 178767, 183263, 188836, 199417, 215506, 219565, 249091, 255355, 277147, 283479, 292013, 295305, 432464, 438448, 503540, 515512, 516173, 535543, 620813, 638263, 640043, 654257 |
| 85030.0 | Geodermatophilaceae | 1382, 6461, 7517, 29713, 51830, 184471, 198403, 198589, 198952, 199152, 199500, 219482, 219514, 247602, 252403, 252630, 253027, 253175, 253272, 262362, 292507, 297607, 297781, 297836, 298197, 305560, 321490, 468828, 470470, 541969 |
| 91061.22 | Bacilli | 1279, 1651, 2895, 19474, 30147, 31546, 44136, 50929, 51658, 51878, 76382, 78148, 178622, 184128, 188246, 198714, 242949, 254924, 257055, 265711, 270219, 285120, 287101, 295257, 298347, 326671, 429595, 509424, 616608, 678895 |
| 91347.1 | Enterobacterales | 260, 815, 1123, 2084, 2529, 6859, 7841, 7966, 8148, 8216, 11834, 13568, 19025, 29462, 37088, 53401, 59419, 196435, 216612, 217182, 229786, 242222, 242496, 242642, 280899, 288794, 295719, 429602, 461443, 516181 |
| 92860.0 | Pleosporales | 1471, 16935, 16936, 53841, 77535, 193144, 266453, 676310, 756921, 758155, 799324, 799424, 799757, 799770, 800035, 800038, 800040, 800096, 813461, 1140562 |
| 92860.1 | Pleosporales | 2668, 76837, 77069, 77206, 193240, 240883, 596902, 651452, 700596, 730381, 799291, 799764, 799771, 799777, 800036, 800082, 800094, 801428, 1056012, 1140510 |
| 110618.0 | Nectriaceae | 5206, 9096, 16928, 37904, 91967, 219602, 241587, 266152, 544331, 547976, 558688, 582016, 593570, 622797, 730481, 737391, 750502, 813427, 1152087, 1152130 |
| 110618.1 | Nectriaceae | 7978, 16800, 30663, 287544, 288168, 532311, 536711, 546213, 554235, 567885, 569620, 604273, 626568, 694050, 717696, 720189, 736672, 738349, 782241, 813383 |
| 117743.1 | Flavobacteriia | 23541, 50827, 59672, 184231, 186230, 192360, 198379, 199439, 199595, 216840, 236641, 237310, 239068, 248536, 248840, 252687, 261828, 284892, 287527, 289734, 295215, 295468, 307602, 321094, 321938, 484164, 598120, 646211, 652681, 1139369 |
| 117747.1 | Sphingobacteriia | 28992, 31802, 186720, 192756, 192980, 198039, 198850, 215002, 236893, 240912, 244168, 244301, 245627, 260823, 283433, 284120, 286701, 291117, 293331, 294616, 295620, 307244, 326921, 451233, 472777, 505355, 515877, 556237, 614460, 646821 |
| 119045.0 | Methylobacteriaceae | 8207, 9081, 28825, 35489, 52050, 52062, 53043, 199229, 199569, 245444, 245926, 254925, 286997, 295122, 321085, 321258, 321605, 321750, 325614, 403266, 410984, 436483, 441523, 480288, 519439, 564885, 572984, 599312, 605848, 662825 |
| 119060.1 | Burkholderiaceae | 231, 605, 3511, 5045, 8815, 10322, 27020, 31838, 33618, 53474, 54717, 75613, 93802, 185151, 197215, 198866, 235347, 237381, 243719, 270342, 280434, 294982, 295382, 325273, 414233, 494363, 532440, 614287, 655523, 672934 |
| 135613.1 | Chromatiales | 1844, 1962, 2964, 5459, 9102, 17881, 19442, 31631, 34410, 34895, 66624, 189462, 192342, 199496, 199657, 218890, 223759, 232638, 243679, 245474, 252707, 276260, 295707, 295717, 297890, 325372, 426424, 433788, 438795, 548632 |
| 135614.1 | Xanthomonadales | 1010, 4210, 29391, 29708, 33067, 50940, 51386, 51430, 182005, 188464, 198575, 198725, 199420, 245812, 251842, 253740, 254438, 274358, 291286, 291562, 308707, 315167, 315271, 315891, 319980, 462066, 543279, 555859, 639274, 1139971 |
| 135619.1 | Oceanospirillales | 238, 239, 1062, 6286, 14463, 28073, 78070, 182350, 185639, 186895, 187327, 190064, 198500, 198641, 199677, 244044, 254326, 257039, 267342, 276016, 283087, 287023, 317839, 327197, 518892, 547614, 565262, 595332, 599578, 1058168 |
| 135620.0 | Oceanospirillaceae | 11866, 27318, 32749, 33747, 54058, 78070, 92544, 92627, 186895, 191418, 199058, 202440, 236745, 242469, 242999, 250744, 253769, 256542, 282818, 283087, 324760, 325302, 502243, 538931, 565262, 595663, 599578, 628710, 640333, 1150830 |
| 135621.1 | Pseudomonadaceae | 2515, 15503, 16560, 29493, 30063, 32531, 33462, 51864, 56648, 196445, 198706, 199524, 232228, 243063, 243207, 243232, 243299, 243488, 243778, 243924, 244119, 251762, 270661, 278332, 294575, 306753, 315235, 319627, 327522, 509591 |
| 135622.1 | Alteromonadales | 1558, 1982, 2608, 6327, 6334, 6843, 14461, 16487, 19276, 33452, 53586, 70299, 194841, 219285, 244441, 247426, 256561, 286482, 294826, 297642, 304912, 307702, 316850, 327086, 481517, 502608, 601768, 606935, 623842, 682739 |
| 135623.0 | Vibrionales | 593, 2493, 2943, 3604, 16562, 16570, 16895, 29994, 31672, 32303, 33633, 37515, 51221, 92876, 94070, 94165, 94936, 184608, 188276, 193432, 198854, 215148, 219336, 235828, 281112, 462621, 464262, 533102, 535589, 1155586 |
| 147541.0 | Dothideomycetes | 8062, 11761, 16931, 16933, 75602, 183809, 194280, 225277, 266453, 676310, 700596, 756921, 785799, 799324, 799757, 799777, 800035, 800040, 801428, 809789 |
| 147541.1 | Dothideomycetes | 30672, 30706, 33647, 77206, 215127, 240883, 243723, 298493, 325902, 504637, 758155, 799424, 799436, 799750, 799767, 800092, 800094, 800235, 813461, 1056012 |

**Supplementary Table 3.** (Continued.)

| Set No. | Taxon name | Proteome IDs |
| --- | --- | --- |
| 147541.2 | Dothideomycetes | 1067, 1471, 2668, 16934, 16936, 53259, 53841, 77069, 237631, 243797, 308549, 310066, 316270, 660729, 799423, 799429, 799439, 799441, 799770, 1140510 |
| 147541.3 | Dothideomycetes | 27730, 70133, 76837, 192596, 309340, 504636, 596902, 651452, 714618, 730381, 799437, 799438, 799537, 799771, 800036, 800038, 800039, 800041, 800082, 1140562 |
| 147545.0 | Eurotiomycetes | 1261, 2530, 7304, 19471, 24533, 42958, 53328, 53599, 184304, 184546, 191612, 224080, 246702, 248961, 249363, 325672, 606974, 661057, 1149074, 1150879 |
| 147545.1 | Eurotiomycetes | 560, 2669, 6702, 6706, 19478, 36893, 55045, 70168, 191342, 191408, 214365, 247810, 256690, 286921, 326565, 326950, 1141434, 1147782, 1149163, 1150904 |
| 147545.2 | Eurotiomycetes | 724, 1294, 1745, 2038, 2497, 54342, 191518, 226031, 234275, 234585, 247233, 248340, 248349, 326289, 452235, 1147747, 1148299, 1149165, 1150941, 1153618 |
| 147545.3 | Eurotiomycetes | 19473, 27920, 53411, 94569, 184188, 191672, 218381, 223968, 231358, 248423, 249402, 249829, 253729, 326198, 509510, 631181, 637239, 1146351, 1147752, 1154252 |
| 147545.4 | Eurotiomycetes | 8142, 19484, 34291, 37505, 53342, 54302, 177622, 182235, 184300, 184499, 186955, 215305, 242791, 243015, 319663, 325780, 326532, 326799, 541154, 641853 |
| 147545.5 | Eurotiomycetes | 1631, 2035, 9882, 18001, 19804, 53617, 54383, 54466, 78343, 179179, 184073, 184383, 188318, 215289, 327118, 701341, 1147733, 1149079, 1149954, 1152649 |
| 147545.6 | Eurotiomycetes | 6701, 19376, 30104, 30143, 37696, 54771, 94526, 184356, 190312, 191285, 191691, 234474, 325579, 326268, 326931, 653565, 654913, 1147746, 1147760, 1150942 |
| 147548.0 | Leotiomycetes | 1798, 6753, 15441, 16922, 178129, 184330, 235371, 241818, 242519, 256645, 283383, 286134, 297299, 297527, 326757, 531561, 624404, 696280, 701801, 1152300 |
| 147550.0 | Sordariomycetes | 5206, 5426, 7322, 8782, 11096, 37136, 76580, 78544, 237481, 242955, 275385, 319160, 326340, 544331, 558688, 694050, 699042, 717696, 720189, 813383 |
| 147550.1 | Sordariomycetes | 1197, 9058, 9096, 16928, 27238, 36947, 54544, 70121, 76744, 194361, 286045, 288168, 532311, 605986, 654918, 722485, 749293, 756346, 1140502, 1152049 |
| 147550.2 | Sordariomycetes | 2762, 6039, 7115, 8066, 9097, 18087, 30651, 54821, 193689, 233524, 241690, 285146, 305883, 481858, 515153, 536711, 547976, 696573, 738349, 1152087 |
| 147550.3 | Sordariomycetes | 2499, 34112, 76552, 78397, 78576, 91967, 222788, 236621, 240493, 241462, 243732, 266152, 283895, 297716, 557566, 567885, 574317, 635477, 827724, 829364 |
| 147550.4 | Sordariomycetes | 1610, 30753, 70720, 76874, 94444, 226192, 235728, 236664, 241587, 265631, 266234, 287144, 310108, 315783, 604273, 622797, 626568, 736672, 781932, 1152130 |
| 147550.5 | Sordariomycetes | 1881, 7978, 8513, 14480, 15530, 16800, 78340, 245910, 266272, 277212, 287544, 325492, 434172, 554235, 562929, 569620, 593570, 777438, 811619, 824596 |
| 147550.6 | Sordariomycetes | 8181, 8984, 16801, 16923, 28524, 37904, 54516, 76584, 76881, 295703, 517252, 582016, 639643, 730481, 750502, 782241, 813426, 813427, 813444, 830671 |
| 147550.7 | Sordariomycetes | 1805, 7796, 20467, 27002, 30106, 30663, 30816, 31176, 70501, 176998, 182658, 186583, 241546, 243498, 252748, 284375, 546213, 737391, 748025, 758603 |
| 155619.0 | Agaricomycetes | 27222, 53257, 54097, 54166, 54477, 54485, 54538, 54988, 217199, 217790, 219338, 230002, 283269, 287166, 294933, 623687, 629468, 772434, 807306, 813600 |
| 155619.1 | Agaricomycetes | 17559, 27073, 27195, 27265, 53558, 53712, 53989, 76761, 76871, 77086, 193067, 284706, 313359, 521943, 664032, 736335, 799118, 807469, 1049176, 1142393 |
| 155619.2 | Agaricomycetes | 1861, 7431, 8493, 30669, 53477, 54018, 76154, 92154, 184267, 194127, 218811, 250043, 305948, 307440, 320762, 322482, 365756, 703269, 823399, 1150266 |
| 155619.3 | Agaricomycetes | 6352, 16930, 30671, 54007, 76532, 76727, 77266, 218334, 242287, 292082, 298030, 305067, 308652, 521872, 609673, 714275, 724874, 807342, 807769, 812287 |
| 165695.0 | *Sphingobium* | 1275, 7753, 9887, 13201, 15523, 15524, 18867, 24284, 52232, 56968, 60779, 78304, 78322, 94323, 199142, 218272, 239825, 267239, 279959, 282977, 290958, 290975, 426201, 549617, 552700, 552757, 571950, 576821, 681425, 1138757 |
| 165696.0 | *Novosphingobium* | 4030, 4728, 7515, 9134, 9242, 37998, 50893, 52268, 56630, 58012, 61032, 199304, 220675, 232587, 236327, 295319, 321172, 323719, 426325, 431236, 465810, 522081, 548867, 551327, 555448, 562395, 566813, 608154, 617634, 648075 |
| 171549.1 | Bacteroidales | 3089, 3167, 5436, 5580, 5697, 15993, 17831, 18054, 18057, 18062, 18111, 18392, 18727, 23482, 29597, 30116, 30150, 56419, 95517, 177035, 195772, 199452, 216094, 235918, 251835, 297225, 324383, 500961, 555103, 629836 |
| 171552.0 | Prevotellaceae | 2772, 2786, 3112, 3160, 3327, 3460, 3829, 4477, 5141, 5580, 10433, 10460, 10862, 15929, 15993, 16600, 17958, 18116, 18343, 18353, 18359, 18375, 70533, 198673, 198779, 199132, 216094, 216769, 272376, 825483 |
| 186801.13 | Clostridia | 3244, 5561, 8234, 18023, 18124, 29585, 31366, 32483, 35704, 50326, 175744, 182321, 182569, 184342, 184869, 190657, 196526, 198636, 199228, 199394, 242329, 243547, 261080, 271256, 287890, 294919, 503601, 515703, 617951, 657006 |
| 186802.6 | Eubacteriales | 4073, 4754, 8937, 10847, 14108, 17961, 18037, 18082, 18137, 19482, 29585, 29916, 36923, 76603, 78233, 184080, 184447, 186102, 195869, 198597, 220840, 236356, 264338, 287890, 298460, 323166, 474718, 597668, 651482, 661435 |

**Supplementary Table 3.** (Continued.)

| Set No. | Taxon name | Proteome IDs |
|---|---|---|
| 186803.3 | Lachnospiraceae | 4121, 5396, 5561, 8178, 17921, 18002, 18056, 18163, 18690, 95409, 183918, 184038, 186530, 195974, 196053, 198806, 198838, 199418, 199659, 199701, 199786, 242090, 245488, 250003, 260812, 292927, 295711, 652477, 655830, 712157 |
| 186806.0 | Eubacteriaceae | 4754, 5178, 12589, 14176, 17904, 17940, 17954, 17996, 18124, 18312, 18318, 18404, 18411, 18414, 36873, 184251, 189857, 190657, 190814, 196365, 198817, 199228, 199394, 253490, 262191, 284400, 284779, 440004, 616595, 663499 |
| 186817.5 | Bacillaceae | 594, 6316, 11747, 27980, 30401, 54099, 74108, 180199, 193006, 198778, 199066, 199668, 216498, 226441, 247922, 263943, 267430, 270219, 271374, 276349, 297982, 325032, 615181, 637720, 660110, 666543, 678477, 678491, 1084197, 1145069 |
| 186818.0 | Planococcaceae | 6691, 11919, 19062, 30408, 30416, 31496, 31938, 58129, 67683, 70230, 93199, 188184, 217065, 222866, 245938, 251002, 254519, 265725, 271255, 275473, 288623, 295327, 297458, 315753, 316502, 321901, 473875, 622653, 658225, 680589 |
| 186822.1 | Paenibacillaceae | 1877, 3445, 29487, 29512, 36611, 37269, 61660, 77355, 78148, 93309, 93585, 187172, 193834, 195591, 198936, 215509, 215741, 247476, 266482, 272464, 289396, 298246, 316330, 317036, 321157, 478765, 574133, 632125, 650466, 677034 |
| 186826.0 | Lactobacillales | 432, 449, 2707, 5147, 7946, 8129, 12675, 16057, 18559, 32287, 51296, 51450, 51679, 51804, 52013, 70387, 182149, 182508, 187499, 191200, 195918, 198374, 198833, 234775, 242754, 279194, 287857, 288197, 310506, 1139006 |
| 194924.0 | Desulfovibrionaceae | 2194, 2601, 2710, 4662, 6034, 6424, 11724, 14960, 14975, 16587, 91979, 181901, 186323, 186469, 189733, 190027, 198324, 218812, 219215, 266086, 267796, 269883, 438699, 448292, 461162, 469724, 473968, 494245, 503820, 580856 |
| 200643.4 | Bacteroidia | 3167, 3327, 3460, 4156, 4477, 4524, 5697, 10460, 18026, 18121, 18343, 18392, 30103, 30150, 76586, 95517, 177035, 184436, 184509, 191055, 196154, 198779, 199132, 215155, 243574, 249300, 255233, 279562, 282985, 297225 |
| 200644.6 | Flavobacteriales | 1602, 4690, 5638, 5938, 6650, 76715, 93807, 184225, 192393, 195822, 198384, 198792, 199448, 199595, 231960, 232401, 233435, 238805, 247903, 249542, 254305, 261828, 321080, 321954, 443153, 484164, 608754, 651057, 656244, 712080 |
| 200666.1 | Sphingobacteriales | 852, 2774, 28992, 31802, 50543, 62859, 95113, 178216, 189739, 189981, 192756, 199577, 199666, 238642, 244168, 245627, 248198, 284120, 291117, 294616, 295499, 295668, 310477, 320042, 325105, 325329, 521691, 646821, 651668, 1139450 |
| 200940.1 | Thermodesulfobacteriota | 1933, 2420, 2534, 2710, 5778, 6365, 6424, 6583, 8561, 14975, 14977, 16587, 68196, 75994, 182264, 191931, 198771, 199355, 199602, 218812, 269883, 317155, 318307, 324159, 422108, 494220, 525298, 596092, 663722, 1144372 |
| 201174.32 | Actinomycetota | 4830, 14204, 15001, 28870, 34098, 35034, 51640, 95759, 192775, 199323, 221369, 270471, 277858, 292547, 321196, 325988, 326179, 463027, 466517, 467328, 503297, 521922, 540423, 557204, 614239, 617734, 619355, 619453, 654257, 1138997 |
| 203682.1 | Planctomycetota | 1887, 6860, 11885, 11991, 76318, 76398, 187735, 199518, 245802, 280296, 315017, 315440, 315724, 317238, 318017, 318053, 318478, 318741, 318878, 318995, 320496, 322699, 324233, 324479, 324974, 326837, 464378, 551616, 676194, 1155241 |
| 203683.0 | Planctomycetia | 1887, 6860, 76398, 187735, 199518, 245802, 280296, 315017, 315440, 315471, 315700, 316095, 316598, 317421, 317648, 317835, 318017, 318053, 318437, 318538, 318995, 319976, 320496, 320672, 322214, 324974, 464378, 503447, 551616, 1155241 |
| 203691.0 | Spirochaetota | 332, 503, 811, 1408, 1847, 2967, 3571, 5737, 6546, 6852, 7383, 8212, 9222, 9223, 14541, 18680, 42527, 182360, 190423, 253869, 294625, 297855, 298058, 323824, 324209, 515827, 518887, 578697, 587760, 660053 |
| 203692.0 | Spirochaetia | 1807, 2318, 2967, 5737, 6048, 6546, 7254, 7383, 9222, 14541, 14605, 15620, 18680, 182360, 186400, 190395, 190423, 192343, 245133, 294625, 297855, 298058, 323824, 324209, 324638, 515827, 518887, 578697, 587760, 671908 |
| 204432.0 | Terriglobia | 343, 2207, 2432, 6056, 6844, 7113, 182427, 198356, 199024, 236728, 253606, 264702, 269669, 289437, 290253, 292958, 295210, 321820, 513480, 515312, 515460, 515496, 538666, 540989, 568106, 589520, 593892, 647241, 648801, 1059380 |
| 204433.0 | Terriglobales | 343, 2207, 2432, 6056, 6844, 7113, 182427, 198356, 199024, 236728, 253606, 264702, 269669, 279598, 289437, 290253, 292958, 295210, 321820, 513480, 515312, 515460, 515496, 538666, 540989, 568106, 589520, 647241, 648801, 1059380 |
| 204441.3 | Rhodospirillales | 245, 7058, 32356, 179145, 185678, 186308, 216998, 231658, 237344, 239724, 245629, 245765, 247565, 277007, 280346, 295023, 295096, 305654, 315252, 317730, 321405, 321746, 553706, 580654, 581135, 600101, 631034, 677537, 761264, 830341 |
| 204455.3 | Rhodobacterales | 2931, 3635, 6762, 18782, 23430, 27746, 31326, 176562, 184932, 192273, 193409, 198307, 198761, 199180, 199344, 202485, 215377, 238338, 244924, 248916, 295142, 309747, 322545, 434754, 436694, 468591, 477083, 526408, 609121, 639775 |
| 204457.4 | Sphingomonadales | 9134, 11717, 11816, 13072, 18851, 27647, 33202, 37930, 55668, 78322, 195807, 198359, 219494, 236327, 254856, 309389, 320547, 325427, 431236, 445582, 466966, 471147, 471435, 509259, 516148, 522753, 538147, 546200, 575068, 584526 |

**Supplementary Table 3.** (Continued.)

| Set No. | Taxon name | Proteome IDs |
|---|---|---|
| 204458.0 | Caulobacterales | 1364, 1492, 1816, 1868, 2696, 17808, 17812, 17826, 17837, 37838, 50923, 51150, 51182, 51604, 56905, 195546, 228945, 244913, 247763, 249524, 249725, 249842, 317223, 431269, 482044, 548978, 622580, 662572, 663918, 676409 |
| 206351.0 | Neisseriales | 1274, 1424, 2010, 3009, 3019, 4088, 4105, 4207, 17813, 18554, 31390, 36114, 76625, 188659, 198213, 215450, 219669, 230202, 242869, 244173, 254293, 254651, 282438, 310016, 325536, 504844, 509597, 543030, 645257, 653156 |
| 206389.0 | Rhodocyclales | 2186, 6552, 13140, 70186, 186819, 187526, 198607, 215181, 217984, 234247, 241193, 241885, 242205, 244972, 248259, 257259, 295129, 307956, 308430, 318422, 323671, 389128, 463961, 561045, 580043, 587070, 599523, 663379, 663444, 694660 |
| 213115.0 | Desulfovibrionales | 1052, 2194, 2216, 2601, 4662, 6424, 7844, 14960, 14975, 16587, 91979, 95200, 181901, 186323, 189733, 190027, 198324, 198771, 199355, 199602, 218812, 219215, 240513, 267796, 269883, 448292, 461162, 469724, 494245, 503820 |
| 213849.0 | Campylobacterales | 939, 2495, 5085, 5709, 7032, 8633, 8721, 15520, 29920, 34444, 92884, 186074, 194309, 201169, 230780, 249746, 251062, 251135, 251881, 255139, 256421, 256695, 257067, 319322, 326944, 476338, 502820, 503264, 509414, 671852 |
| 216572.1 | Oscillospiraceae | 2145, 3438, 4259, 14129, 18037, 18181, 18315, 18328, 19365, 29585, 196146, 196191, 196537, 199158, 233534, 253465, 264338, 276301, 285063, 301475, 306409, 324781, 469514, 474718, 597668, 610760, 620327, 660861, 661435, 774750 |
| 335929.0 | Erythrobacteraceae | 2995, 16568, 49978, 53070, 53455, 77326, 78263, 92932, 275232, 284395, 309389, 315722, 320547, 325427, 326213, 429229, 431922, 433104, 433652, 439522, 445582, 448199, 460561, 461409, 469159, 469430, 473531, 502110, 546031, 594459 |
| 423349.0 | *Mucilaginibacter* | 2774, 189739, 199072, 199679, 215002, 242687, 244168, 245678, 253209, 260823, 268007, 282759, 286701, 293331, 318733, 320042, 321479, 429232, 503278, 505355, 521691, 538158, 556237, 613193, 619078, 622475, 638732, 646821, 662074, 1139450 |
| 563835.0 | Chitinophagaceae | 3586, 31408, 33121, 77177, 77667, 184420, 192276, 192610, 192796, 198711, 199031, 199537, 217148, 239872, 240572, 240971, 242818, 244450, 249720, 263900, 267223, 279089, 281028, 290204, 305848, 321204, 321513, 321533, 326903, 1155483 |
| 681950.0 | Glomerellaceae | 8782, 11096, 14480, 15530, 20467, 27238, 70121, 76552, 176998, 186583, 295703, 305883, 326340, 434172, 613401, 639643, 654918, 699042, 781932, 830671 |
| 768503.1 | Cytophagia | 493, 2011, 10796, 50454, 59956, 61382, 75267, 75606, 189843, 192266, 193660, 198432, 199437, 199513, 234203, 245468, 251692, 295706, 297549, 297647, 322791, 441336, 480178, 501128, 509425, 515410, 537126, 598820, 603640, 609064 |
| 768507.1 | Cytophagales | 2011, 9309, 10953, 19423, 30789, 50454, 51810, 61382, 75583, 184212, 184609, 192266, 192333, 194873, 198432, 198510, 199306, 245468, 251692, 297549, 305398, 317847, 478546, 501623, 502756, 515410, 603640, 612233, 632594, 664144 |
| 909932.0 | Negativicutes | 1902, 3195, 3277, 3503, 4923, 7093, 9883, 10111, 11124, 14380, 14944, 16614, 29628, 36503, 41641, 95546, 182958, 195529, 198896, 199309, 199689, 215383, 276437, 277811, 284368, 295063, 295188, 320776, 323646, 591941 |
| 1028384.0 | Glomerellales | 8782, 11096, 14480, 15530, 20467, 27238, 70121, 76552, 76584, 176998, 186583, 295703, 305883, 326340, 434172, 613401, 639643, 699042, 781932, 830671 |
| 1131492.0 | Aspergillaceae | 9882, 19804, 70168, 177622, 179179, 184188, 184546, 215289, 218381, 246702, 247233, 248349, 248423, 325579, 326289, 452235, 1146351, 1147760, 1147782, 1149954 |
| 1131492.1 | Aspergillaceae | 2530, 6702, 19376, 34291, 37696, 94569, 184300, 184304, 184383, 186955, 188318, 190312, 191285, 248961, 249402, 325780, 326950, 327118, 631181, 1149163 |
| 1131492.2 | Aspergillaceae | 6701, 30104, 36893, 37505, 42958, 54771, 191408, 231358, 234474, 248340, 256690, 319663, 1147733, 1147746, 1148299, 1149074, 1150879, 1150941, 1150942, 1152649 |
| 1131492.3 | Aspergillaceae | 560, 30143, 184356, 184499, 215305, 234275, 247810, 253729, 326198, 326268, 326532, 326799, 326931, 630445, 637239, 641853, 1147747, 1150904, 1153618, 1154252 |
| 1131492.4 | Aspergillaceae | 724, 6706, 55045, 184073, 191342, 191518, 191612, 191672, 234585, 286921, 325672, 326565, 541154, 653565, 654913, 661057, 701341, 1147752, 1149079, 1149165 |
| 1643682.0 | Geodermatophilales | 1382, 6461, 7517, 29713, 51830, 184471, 198403, 198589, 198857, 198952, 199152, 199500, 219482, 219514, 237752, 252403, 252630, 253027, 253175, 253272, 262362, 292507, 297607, 297781, 298197, 305560, 321490, 468828, 470470, 541969 |
| 1706369.0 | Cellvibrionales | 1036, 1947, 4699, 9080, 29640, 76077, 77184, 188219, 193450, 196143, 234845, 267187, 273643, 275394, 294980, 295554, 319732, 326835, 467441, 528457, 535937, 539350, 559987, 596063, 610558, 652567, 664303, 787472, 1056808, 1069090 |
| 1737404.0 | Tissierellia | 1319, 3280, 3422, 3705, 4191, 4712, 5286, 6094, 17105, 31386, 32409, 37267, 70442, 179042, 180254, 184032, 184114, 184389, 186112, 198828, 245423, 261011, 269544, 284177, 294567, 297454, 298381, 377798, 1108123, 1142078 |
| 1737405.0 | Tissierellales | 1319, 3280, 3422, 3705, 3821, 4191, 4712, 5984, 6094, 13378, 17105, 31386, 32409, 37267, 70442, 179042, 180254, 184032, 184114, 184389, 186112, 198828, 245423, 261011, 294567, 297454, 298381, 377798, 1108123, 1142078 |

**Supplementary Table 3.** (Continued.)

| Set No. | Taxon name | Proteome IDs |
|---|---|---|
| 1775411.0 | Rhodanobacteraceae | 3226, 5234, 29708, 76131, 182005, 184078, 188464, 189228, 198575, 198725, 199420, 199603, 235676, 241074, 245812, 253740, 267077, 270530, 288967, 291822, 294599, 295293, 306317, 307749, 490980, 521199, 543279, 555859, 667269, 1139971 |
| 1822464.0 | *Paraburkholderia* | 1192, 3511, 5045, 30460, 75613, 76852, 93802, 185151, 187012, 198638, 198866, 199120, 199548, 237381, 254875, 263097, 272778, 294982, 325273, 433577, 434209, 494119, 494252, 494255, 494329, 494363, 494365, 655523, 676887, 1139308 |
| 1853228.1 | Chitinophagia | 31408, 77667, 184368, 184420, 185003, 186917, 190166, 190888, 192796, 198757, 199041, 217148, 239872, 240572, 242818, 249547, 253410, 266118, 279089, 281028, 290204, 290545, 292424, 293874, 305848, 321436, 321513, 326903, 476411, 627292 |
| 1853229.1 | Chitinophagales | 3586, 31400, 33121, 77177, 184420, 188534, 190166, 190888, 192276, 192610, 198757, 199041, 199537, 217148, 244333, 244450, 248745, 249720, 266118, 279089, 290204, 293874, 294498, 305848, 321204, 321533, 476411, 569858, 598971, 607559 |
| 1853232.0 | Hymenobacteraceae | 19423, 28590, 30789, 33109, 36458, 59542, 59956, 70672, 182491, 184418, 185924, 189843, 192266, 198432, 198697, 251692, 253919, 262802, 321532, 321926, 322791, 326570, 441336, 478546, 515410, 559626, 563094, 603640, 632594, 664144 |
| 1866885.0 | *Mycolicibacterium* | 757, 4915, 5442, 6057, 6265, 9159, 18763, 28870, 32221, 36176, 57134, 70612, 94243, 178953, 192801, 193484, 193564, 220914, 254978, 279306, 282551, 323700, 325690, 466445, 466517, 466931, 467006, 467193, 467249, 1140293 |
| 1890424.0 | Synechococcales | 788, 1115, 1420, 1422, 1423, 1425, 1430, 1566, 1938, 1961, 2535, 3950, 4972, 5766, 10379, 10385, 10482, 28237, 28591, 76867, 182608, 238762, 240206, 243002, 325996, 515370, 594525, 594557, 607397, 889800 |
| 1913637.0 | Mucoromycota | 14254, 18888, 27586, 53815, 77051, 77315, 93000, 242146, 242381, 247702, 252139, 266673, 646827, 654370, 702732, 707451, 726737, 740926, 764092, 1153678 |
| 1913637.1 | Mucoromycota | 54107, 78512, 78561, 193560, 193648, 234323, 242180, 242254, 263633, 265703, 439903, 603453, 650833, 696485, 716291, 723463, 738325, 748756, 764349, 807716 |
| 2691354.0 | Pirellulales | 1025, 11885, 11991, 76398, 315440, 315471, 316213, 316304, 316598, 316770, 317421, 317429, 317648, 317909, 317977, 318053, 318288, 318437, 318538, 318878, 318995, 319143, 319852, 320176, 321353, 322699, 324479, 325286, 536179, 551616 |
| 2726947.0 | Mycosphaerellales | 8062, 16931, 16932, 16933, 33647, 70133, 75602, 215127, 225277, 237631, 308549, 309340, 310066, 504637, 660729, 756132, 799537, 799539, 799767, 1056384 |
| 2762318.0 | Weeksellaceae | 6051, 6085, 6276, 8641, 28641, 35900, 92708, 95601, 182761, 184335, 188947, 191112, 192393, 198517, 198931, 215196, 238042, 252172, 270538, 273471, 279525, 294419, 309120, 464318, 552241, 553459, 608754, 610746, 694480, 1152599 |
| 2854170.3 | Roseobacteraceae | 2944, 5713, 19593, 26249, 31171, 31326, 48908, 184221, 192455, 193409, 193570, 198634, 198728, 198767, 198994, 199026, 199283, 199550, 199628, 202922, 203589, 244912, 252940, 274386, 284407, 293145, 305887, 325526, 327025, 503308 |
| 2887326.0 | Moraxellales | 430, 546, 5740, 13117, 13148, 14568, 18415, 18418, 23785, 23788, 23795, 35860, 64939, 76238, 92616, 92671, 93391, 185895, 187495, 188357, 189800, 190435, 196536, 242317, 243900, 244223, 245977, 253940, 255230, 256774 |
| 2975441.0 | Sphaerotilaceae | 366, 1693, 7883, 53102, 56576, 60699, 193427, 197446, 197468, 199246, 238605, 239406, 247811, 274884, 288178, 288587, 293433, 295110, 295361, 301751, 323522, 430120, 484255, 518288, 529637, 554837, 574067, 594526, 620139, 678374 |
| 3028117.0 | Cyanophyceae | 268, 1115, 1191, 1203, 1566, 5766, 10383, 10384, 10386, 10390, 10477, 10483, 16960, 28591, 29738, 182608, 215660, 218785, 243002, 252107, 271624, 282767, 503129, 596184, 607678, 619131, 627856, 627988, 632766, 645501 |
| 3031449.0 | Desulfovibrionia | 1052, 2194, 2216, 2601, 4662, 5496, 6424, 7844, 11724, 14975, 16587, 91979, 95200, 181901, 186323, 186469, 189733, 190027, 198771, 199355, 199602, 218812, 240513, 267796, 269883, 438699, 448292, 461162, 469724, 473968 |
| 3031852.0 | Epsilonproteobacteria | 422, 760, 1136, 2222, 2407, 5085, 5709, 6431, 7032, 7727, 7803, 8387, 8633, 14539, 18688, 29313, 194309, 229442, 230780, 251062, 251881, 254920, 256421, 256695, 257067, 272365, 451105, 476338, 503264, 509414 |
| 3082720.0 | Peptostreptococcales | 269, 3178, 3244, 5244, 17980, 19591, 31189, 34407, 93352, 184082, 184465, 190140, 190285, 198304, 198718, 199208, 199230, 199568, 215694, 216024, 242520, 243255, 243494, 245695, 284841, 287601, 294919, 295500, 295504, 503601 |
| 3085636.3 | Lachnospirales | 370, 1476, 17909, 18002, 37233, 95380, 95409, 183918, 184301, 198508, 198838, 199315, 199800, 199802, 199820, 236497, 245488, 280696, 285302, 290106, 292927, 295711, 298653, 460412, 464314, 483018, 543642, 623269, 677305, 712157 |

**Supplementary Table 4.** List of 12 structural core genes defined from 166 species spanning the tree of life. For each core gene, representative identifier and functional description annotated by UniProt are described, as well as overlapping status against OrthoFinder.

| Identifier | Functional desciption | OrthoFinder |
|---|---|---|
| A0A196SAG3 | Large ribosomal subunit protein uL15 | Not found |
| A0A2S8SPM9 | Amidophosphoribosyltransferase | Found |
| A0A3G2TFK5 | Large ribosomal subunit protein uL13 | Found |
| A0A7G8BI38 | Small ribosomal subunit protein uS9 | Not found |
| A0A8K0DZM5 | Protein translocase subunit SecY | Not found |
| A0A9D3NI02 | Adenylosuccinate synthetase | Found |
| A0A9P6U5R7 | Tyrosine--tRNA ligase | Found |
| A0A9Q1C2Q7 | Large ribosomal subunit protein uL1 | Not found |
| B4CUY6 | Large ribosomal subunit protein uL5 | Not found |
| B8C5S9 | CTP synthase | Found |
| E3MII4 | Aspartyl/glutamyl-tRNA amidotransferase subunit B | Found |
| Q0BYC9 | Large ribosomal subunit protein uL6 | Not found |
